# Matrin3 mediates differentiation through stabilizing chromatin accessibility and chromatin loop-domain interactions, and YY1 mediated enhancer-promoter interactions

**DOI:** 10.1101/2023.03.30.534959

**Authors:** Tianxin Liu, Qian Zhu, Kai Yan, Trevor Bingham, Hye Ji Cha, Stuti Mehta, Thorsten M Schlaeger, Guo-Cheng Yuan, Stuart H Orkin

## Abstract

Although emerging evidence indicates that alterations in proteins within nuclear compartments elicit changes in chromosomal architecture and differentiation, the underlying mechanisms are not well understood. Here we investigate the direct role of the abundant nuclear complex protein Matrin3 (Matr3) in chromatin architecture and development in the context of myogenesis. Using an acute targeted protein degradation platform (dTAG-Matr3), we reveal the dynamics of development-related chromatin reorganization. Upon acute depletion of Matr3, gains in chromatin accessibility and MyoD binding were observed, prior to widespread loss in the steady-state Matr3-knockout. These initial changes correlated with gene expression changes later in development. High-throughput chromosome conformation capture (Hi-C) experiments revealed substantial chromatin loop rearrangements soon after Matr3 depletion. Notably, YY1 binding was detected in close proximity to enhancer and promoter regions, accompanied by the emergence of novel YY1-mediated enhancer-promoter loops, which occurred concurrently with changes in histone modifications and chromatin-level binding patterns. Overall, our results suggest that Matr3 mediates differentiation through stabilizing chromatin accessibility and chromatin loop-domain interactions, and highlight a conserved and direct role for Matr3 in maintenance of chromosomal architecture.

## Introduction

Nuclear proteins in eukaryotes comprise non-chromatin microgranular, ribonucleoproteins, connecting the nuclear membrane to intranuclear components. Acting as scaffold for attachment of chromatin, the nuclear protein complex serves to support and organize chromatin architecture. Together with lamins and heterogeneous nuclear ribonucleoproteins, Matrin3 (Matr3) is the major constituent of the inner nuclear proteins (Nakayasu et al., 1991, Engelke et al., 2014).

Attention in the literature has focused primarily on the involvement of Matr3 in pre-messenger RNA (mRNA) splicing, mRNA stability, and DNA damage repair and replication (Coelho et al., 2015, Salton et al., 2010, Salton et al., 2011). Mutations in Matr3 contribute to a subset of familial amyotrophic lateral sclerosis (ALS) (Johnson et al., 2014, Boehringer et al., 2017). Matr3 has also been implicated in developmental processes, as its knockout is embryonic lethal at or before the E8.5 neural-fold stage (Quintero-Rivera et al., 2015), and its deficiency promotes neural stem differentiation (Niimori-Kita et al., 2018). Recently, we reported that loss of Matr3 in embryonic stem cells and erythroid precursors elicited changes indicative of accelerated differentiation, and was associated with altered chromatin organization (Cha et al.,2021).

To explore how Matr3 participates in chromatin organization more broadly, here we have investigated its loss in the context of an established and tractable differentiation system, the myoblast cell line, C2C12 (Yaffe and Saxel, 1977, Doynova et al., 2017, Bi et al., 2017, Zhang et al., 2017). Studying an abundant nuclear component, such as Matr3, presents challenges, as analysis of a cellular phenotype following gene knockout and accompanying compensatory changes may confound interpretation of its primary roles. To dissect the role of Matr3 in chromatin organization more directly, we have employed targeted protein degradation (TPD). Acute depletion of Matr3 by TPD afforded the opportunity to interrogate chromatin accessibility, transcription (TF) bind, and chromatin organization in a temporal fashion. Our findings identify a distinct role for Matr3 in stabilizing chromatin architecture during cellular differentiation, further establishing a critical requirement beyond its more commonly appreciated involvement in RNA metabolism.

## Results

### Matr3 loss leads to defects in myogenesis

We first evaluated the effect of *Matr3* loss on myogenesis within the context of differentiation of C2C12 cells. CRISPR/Cas9 editing was used to delete the entire *Matr3* gene body (Supplemental Figure 1a-c). Upon analysis of individual C2C12 knockout clones, we observed clonal variation in growth and differentiation (Supplemental Figure 1d). To circumvent clonal effects, we generated cells depleted of Matr3 in bulk using a single guide delivered as RNP (Figure 1a, Supplemental Figure 2). The Matr3 level was reduced by ∼90% in the bulk population (Figure 1b). Upon differentiation, myotubes depleted of Matr3 exhibited more branches than wildtype, reflective of hypertrophy (Figure 1c). This phenotype mimics that of Duchenne muscular dystrophy (DMD), which is characterized by a high percentage of branched myofibers that are vulnerable to contraction-induced injury (Duddy et al., 2015).

**Figure 1.**
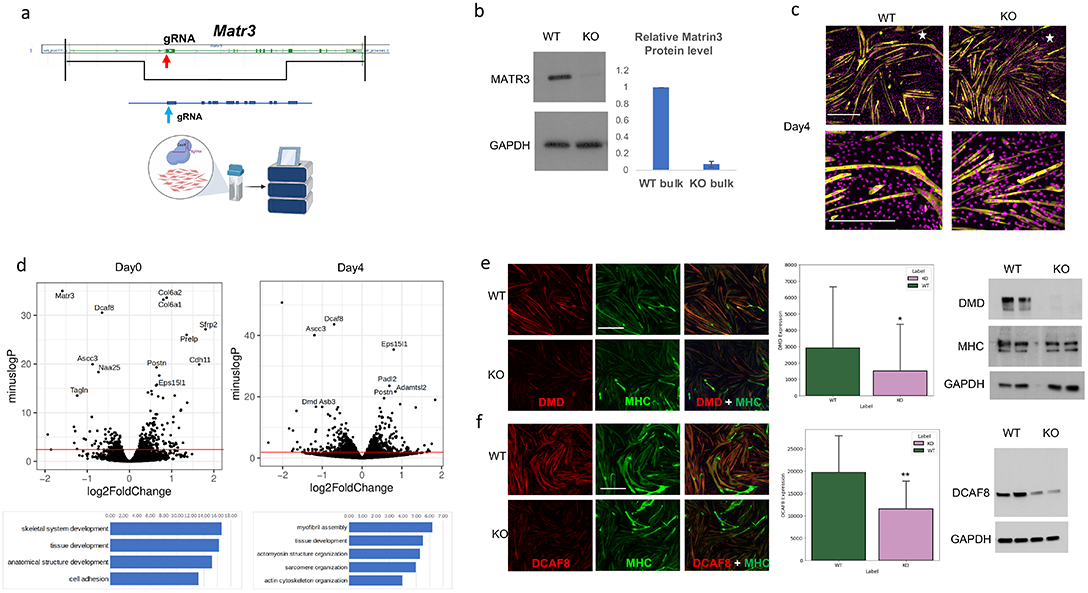
Matr3 loss leads to defects in myogenesis. (a) Generation of Matr3 knockout (KO) bulk using Cas9/RNP transfection (see methods). (b) Matr3 protein level was reduced significantly in Matr3KO bulk. MATR3 expression was assessed in wildtype and Matr3 KO C2C12 cells by Western blot. Representative of the Western blots and the quantification of 3 biological replicates. (c) Myotubes in Matr3KO exhibited hypertrophy phenotype. C2C12 Matr3 KO and wildtype were differentiated for 4 days, and immunostained with DAPI (purple) and Myosin heavy chain (MHC, yellow). Myotubes in Matr3KO (right) had more short branches compared with wildtype (left). The bottom images were magnified views from the regions marked with stars in upper images. All the images were taken from the same regions in each replicate using Yokogawa CV7000 microscope with the same setting. Scale bar, 500um. (d) Knockout of Matr3 contributed to differential gene expression. RNA-seq of cells at myoblast (Day0) and myotube (Day4) stages. Expression changes were measured by KO-WT, p<0.05 for the red line that denotes significant DE genes. DE genes were associated with skeletal muscle development (GO term, bottom panel). (e) Duchenne muscular dystrophy gene (DMD) expression was reduced in Matr3 KO myotubes. C2C12 Matr3 KO and wildtype were differentiated 4 days, and immunostained with DMD (red) and MHC (green). Scale bar, 500um. Signal intensity was quantified based on 3 biological replicates. Protein level of DMD was also confirmed by Western blots (right panel). (f) DCAF8 expression was reduced in Matr3 KO myotubes. C2C12 Matr3 KO and wildtype were differentiated 4 days, and immunostained with DCAF8 (red) and MHC (green). Scale bar, 500um. Signal intensity was quantified from 3 biological replicates. Protein level of DCAF8 was also confirmed by Western blot (right panel).

To assess the impact of Matr3 loss on gene expression, we performed RNA-seq of cells at myoblast (Day 0) and myotube (Day 4) stages. Changes in gene expression were observed, as reflected in enrichment of gene sets associated with skeletal muscle development (Figure 1d). Among differential gene sets, Duchenne muscular dystrophy gene (DMD) expression and DCAF8 were reduced, which was confirmed by protein analyses (Figure 1e-f). Thus, depletion of Matr3 leads to aberrant muscle differentiation.

### Acute depletion of Matr3 contributes to few changes in nascent RNA production

To identify more immediate consequences of Matr3 loss on gene expression and chromatin organization, we employed PROTAC-mediated TPD using the dTAG platform (Nabet et al., 2018, Mehta et al. 2023). Using CRISPR/Cas9- mediated editing, facilitated by AAV template DNA, we engineered C2C12 cells harboring variant FKBP introduced in-frame at the N-terminus of the *Matr3* gene (Figure 2a). Leveraging high-efficiency gene editing in bulk populations, we avoided clonal variation (Figure 2b and Supplemental Figure 3a-b). Following exposure to PROTAC dTAG47, Matr3 protein was depleted within 4 hrs. (Figure 2c, and Supplemental Figure 3c-d).

**Figure 2.**
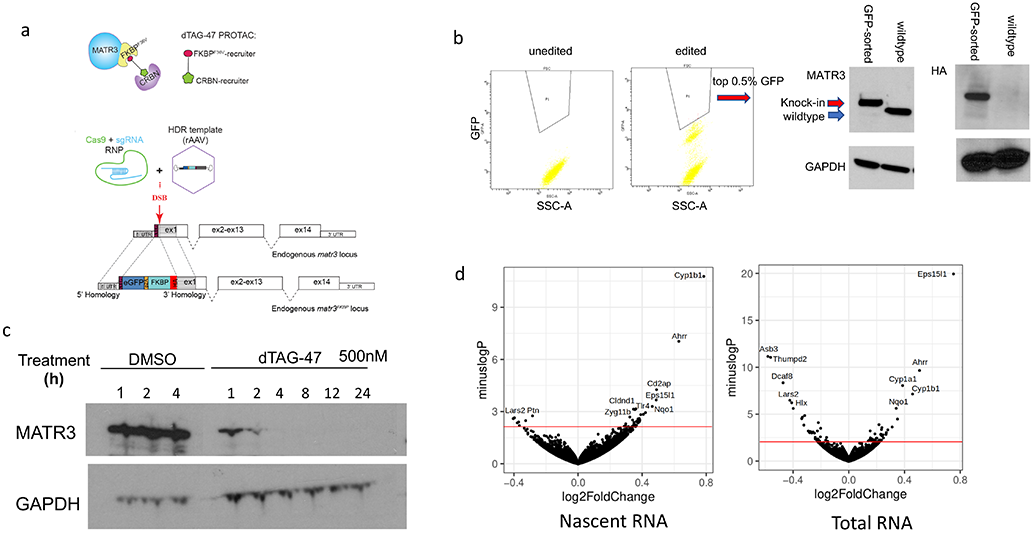
dTAG-Matr3 could be degraded within 4hrs., and acute depletion of Matr3 contributes to few changes in nascent RNA production. (a) Strategy to knockin FKBP ^F36v^ at the 5’ end of Matr3. The dTAG-47 PROTAC recruits the CRBN E3 ligase complex to FKBP ^F36v^-Matr3. (b) High-efficient knockin bulk FKBP ^F36v^-Matr3 were sorted by GFP and confirmed by Western blot. Cells enriched in the top 0.5% highest GFP signal were sorted (GFP-sorted). FKBP ^F36v^-Matr3 (refer as dTAG-Matr3) bands were enriched in the GFP-sorted bulk, and tagging efficiency was also confirmed by HA antibody. (c) Matr3 protein was depleted within 4 hrs. upon dTAG47 exposure. Western blots of dTAG-Matr3 upon dTAG47 (500nM) treatment in a time course. (d) Nascent RNA and total RNA were quantified by SLAM-seq upon 4 hrs. Matr3 depletion. (volcano plots, p<0.05 for red line that denotes significant DE genes).

To identify early transcriptional changes upon Matr3 loss, we performed SLAM-seq at 4 hrs. post dTAG47 exposure and quantified nascent RNA transcription (Herzog et al., 2017). We observed only ∼8 significantly differentially expressed (DE) genes (log_2_FC > 0.25) (Figure 2d). Expanding the DE gene list further, we observed that genes at the top of the DE gene list were highly similar to the 24 hrs. RNA-seq DE gene list (e.g. Eps15l1, Dcaf8, Asb3, Thumpd2, Naa25) (Supplemental Table1), suggesting that critical DE genes are captured by the acute depletion system. Thus, acute depletion of Matr3 (4hrs.) elicited very limited changes in gene expression.

### Acute depletion of Matr3 results in early gains in chromatin accessibility and MyoD binding

Because limited gene expression changes often belie more extensive alterations in the chromatin organization, we next tracked chromatin accessibility upon loss of Matr3 (Buenrostro et al., 2015). In the Matr3 KO system, we observed predominantly loss of chromatin accessibility (Figure 3a). To dissect these changes in detail, we assessed chromatin accessibility upon acute depletion of Matr3. We observed both gains and loss of chromatin accessibility at an early time (4hrs.), which was followed by a trend to greater loss at 8 hrs., and predominant loss thereafter (steady-state). We further assessed the effects of prolonged Matr3 loss on myogenesis by examining chromatin occupancy of the master regulator MyoD by CUT&RUN (Meers et al., 2019, Zhu et al., 2019). We detected a similar trend to that observed for overall chromatin accessibility. Increased MyoD binding was seen at the early time point followed by gradual loss of MyoD binding (Figure 3b). Based on these findings, we surmise that depletion of Matr3 directly initiates chromatin changes characterized by increased accessibility and MyoD occupancy.

**Figure 3.**
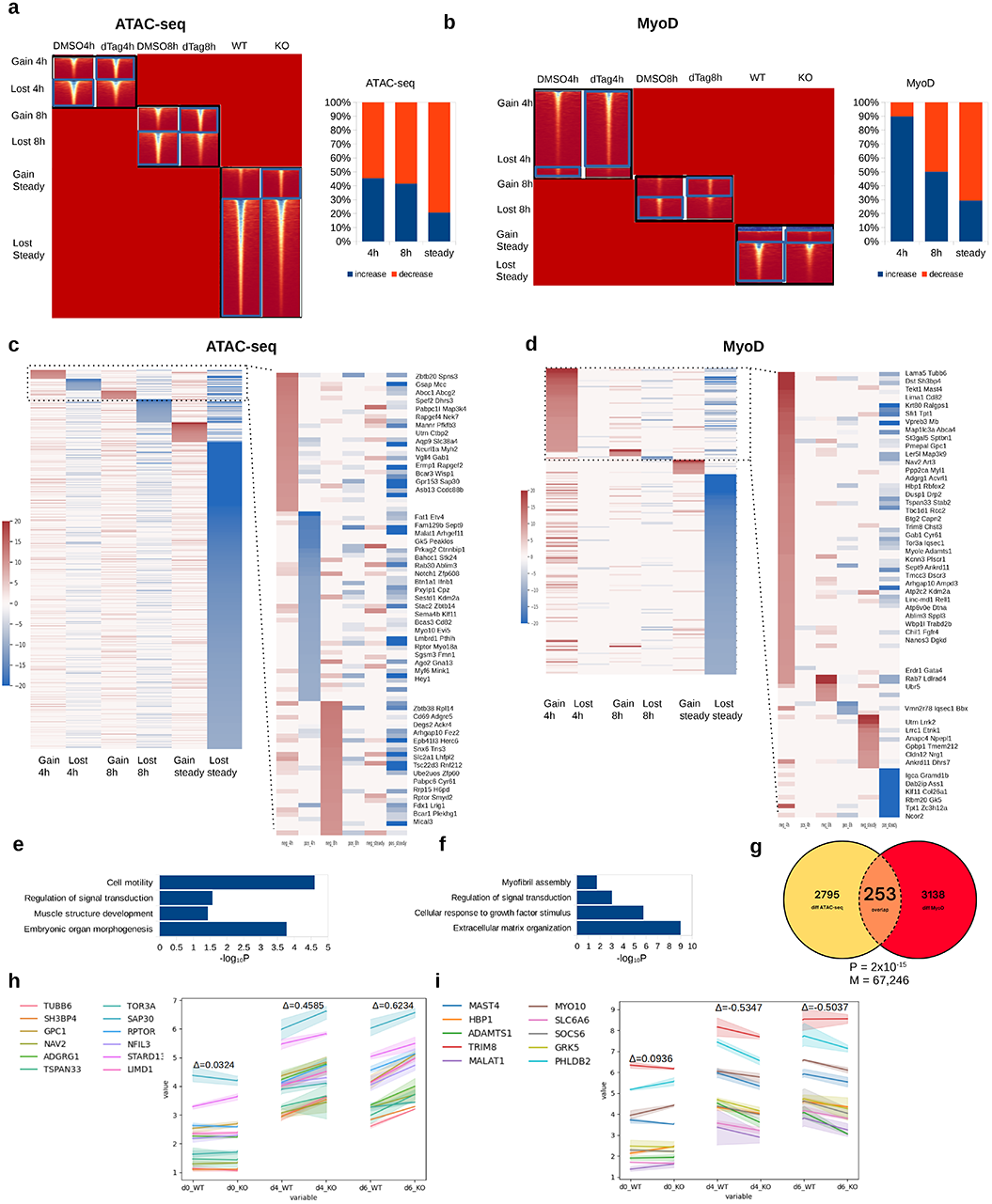
Matr3 depletion alters chromatin accessibility and MyoD binding, which foreshadow later gene expression changes. (a) Heatmap showing differential chromatin accessibility (ATAC-seq) at 4 hrs., 8 hrs. post Matr3 depletion and steady state (Matr3 KO). Bar chart: percentage of peak increased and decreased per time point. (b) Same panels with (a) for MyoD binding (CUT&RUN). Gains trended down over time while losses increased. (c) Genes mapped by differentially chromatin accessible peaks per time point (4hrs., 8hrs., steady), illustrating the magnitude and number of changes. Accessibility was primarily lost at steady state. (d) Genes mapped by differential MyoD binding. Particularly striking were MyoD binding gains at 4hrs. and loss at steady state. (e) GO biological process enrichment on differentially ATAC-seq mapped genes (4hrs.). (f) GO enrichment on differential MyoD bound genes (4hrs.). (g) Overlap of differential ATAC and MyoD bound regions (4hrs.). P: P-value. M: number of co-bound peaks. Number in circle: # of differential peaks. (h-i) Early differentially accessible and MyoD bound genes elicited expression changes later in development (days 4 and 6). Genes were selected from heatmap in panels c and d as having MyoD 4hrs. binding gain or ATAC 4hrs. gain/loss. Expression changes (measured by Δ=KO-WT) were followed during development. Expression changes were not significant at day 0 (steady state), but evident at days 4, and 6. Upregulated genes in Matr3 downregulated genes exhibited positive Δ (h), downregulated genes in Matr3KO show negative Δ (i)

Upon mapping the differentially accessible regions to the genes within 25 kb of the transcription start sites (TSS) (Figure 3c), we observed that loci with increased accessibility included MAPK kinase Map3k4, guanine nucleotide exchange factor Rapgef4, transcription factor Ctbp2, and skeletal muscle target Myh. Gene set enrichment analysis by DAVID (Sherman et al., 2022) and SEEK (Zhu et al., 2015) of differentially accessible genes revealed embryonic organ morphogenesis (P=1.7e-4), muscle structure development (P=3.7e-2), regulation of signal transduction (P=2.7e-2), cell motility/cell migration at 4 hrs. following Matr3 depletion (up-regulated genes) (Figure 3e).

We next asked if early changes in chromatin accessibility could be attributed to redistribution of MyoD occupancy. We mapped differential MyoD occupancy loci to nearby genes (Figure 3d). MyoD differential binding regions were enriched in genes in extracellular matrix organization (P=1e-9), cellular response to growth factor stimulus (P=1.7e-6), myofibril assembly, and regulation of signal transduction (P=9e-4) (Figure 3f). Upon analyzing the overlap between differential ATAC-seq and differential MyoD binding, we observed a total overlap of 253 differential peaks, which represents a highly significant portion of total differential ATAC-seq and MyoD peaks (P=2e-15) (Figure 3g), suggesting that Matr3 depletion simultaneously perturbs open chromatin regions and MyoD binding. Collectively, these results indicate that acute depletion of Matr3 directly initiates changes in chromatin accessibility and MyoD binding at selective sites that are related to signal transduction and development.

### Early perturbed MyoD and chromatin accessibility sites in Matr3 TPD elicit gene expression changes at later developmental stages

Although acute depletion of Matr3 affected differential chromatin accessibility and MyoD chromatin occupancy, few changes in gene expression were observed at early times (Figure 2d). Thus, we hypothesized that early chromatin changes lead to subsequent gene expression changes later in muscle development. We tracked the expression of genes that are downstream of sites with differential chromatin accessibility and MyoD occupancy upon depletion of Matr3 for 4 hrs.. To avoid potential effects of prolonged dTAG47 treatment on cell viability and/or differentiation, we examined gene expression in bulk Matr3 KO cells at different days of differentiation (Figure 3h-i, h for upregulated in Matr3KO, i for downregulated in Matr3KO). Genes nearby perturbed MyoD/accessibility sites exhibited significant gene expression differences in 4 and 6 day differentiated Matr3 KO bulk cells (KO vs WT). No appreciable changes in expression were observed at day 0 of differentiation. Using a large-scale public data-mining approach (Zhu et al., 2015), we deduced a general list of coordinated genes that were co-expressed with genes encoding the Matr3-complex (Pandya-Jones et al., 2021) (Supplemental Figure 4). These co-expressed genes (n=500), representing Matr3-coordinated genes, were most significantly altered in gene expression in day 6 differentiated Matr3 KO cells, indicating that targets of a Matr3-protein complex were altered at a later time by Matr3 loss. Overall, earlier changes in chromatin accessibility and MyoD differential occupancy consequent to Matr3 depletion foreshadowed later developmental gene expression changes.

### Matr3 depletion maintains A/B compartments but rearranges loop domain interactions at a sub-compartment level

Consistent with prior studies of ES and erythroid cells (Cha et al., 2021), depletion of Matr3 (Matr3 KO bulk) in C2C12 cells was associated with changes in CTCF and reduced overall cohesin (assessed by Rad21) occupancy (Supplemental Figure 5a). Notably, changes in CTCF bindings and reduced Rad21 occupancy were observed as early as 4hrs. post depletion of Matr3 (Figure 4a). These findings extend our prior observations by implicating direct involvement of Matr3 in maintaining chromatin organization. To probe this aspect more fully, we performed high-throughput chromosome conformation capture (Hi-C) experiments upon acute depletion of Matr3 (4hrs. dTAG47) (Lieberman-Aiden et al., 2009). No changes were observed in A/B compartment switching and interaction (Supplemental Figure 5b-c). Instead, we detected extensive rearrangement of chromatin loops (Figure 4b, highlighted by circles). We then mapped the rearranged loops on all chromosomes (Figure 4c). Rearranged loops were concentrated in specific genomic, notably within Chr7, Chr14 and ChrX (Figure 4d). As an example, we highlight Tgfb1-Ltbp4 loop rearrangements, which were ranked among the most re-arranged regions on chromosome 7 (Figure 4e). Both gained and lost loops were adjacently located at the intersection point (see circle in Figure 4f). Together, these findings indicate that Matr3 loss extensively perturbs chromatin loop formation.

**Figure 4.**
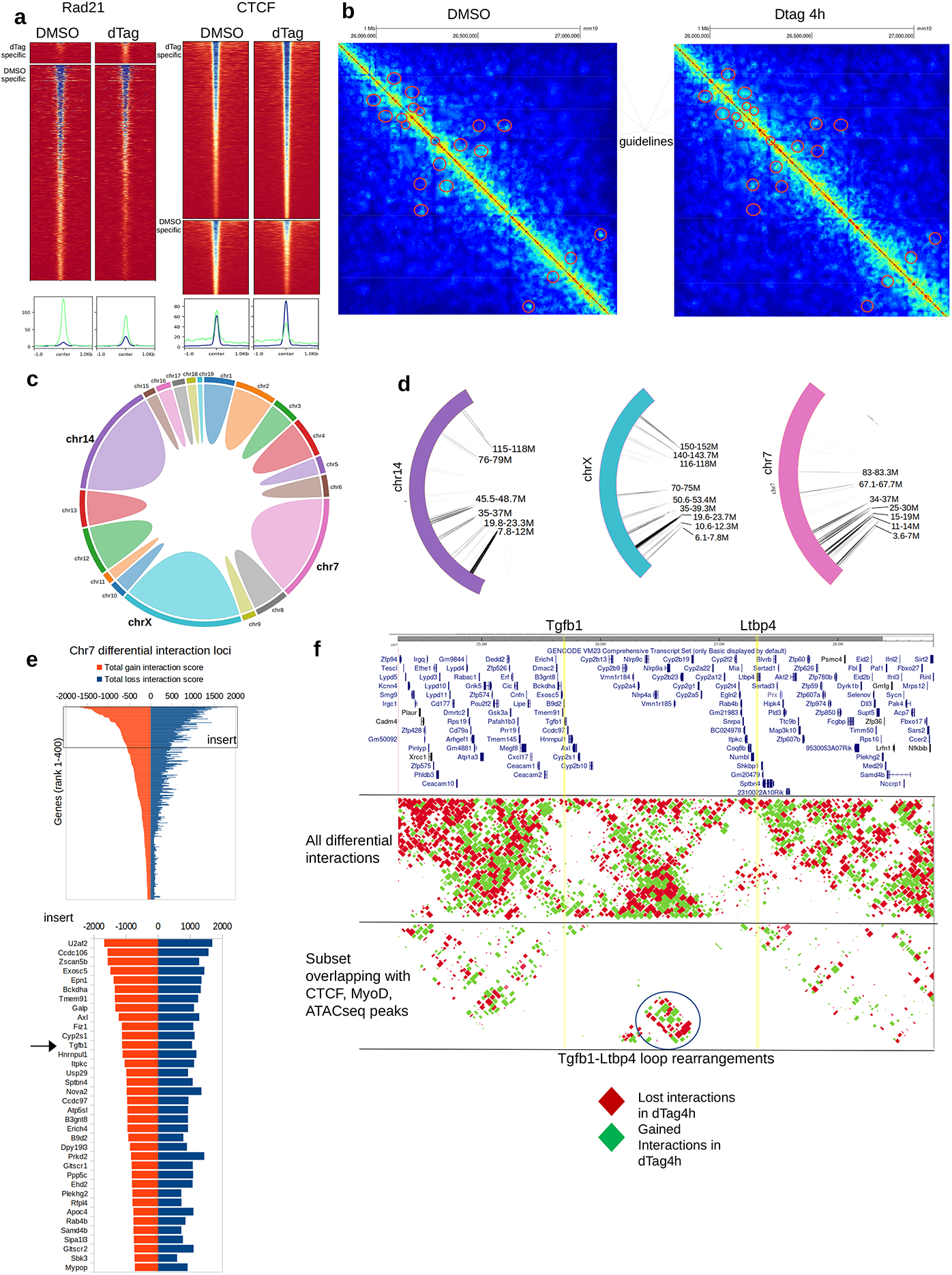
Matr3 depletion extensively rearranges loop domain interactions. (a) Substantial reduction of Rad21 binding 4hrs. following Matr3 depletion. CTCF occupancy was marked by both gains and losses. The histograms below summarize the binding intensities (green: lost peaks, blue: gained peaks). (b) Example of loop rearrangements derived from Hi-C experiments that characterized the effect on chromatin structure at fragment resolution (on average 5kb). Red circles denote sites of differential interactions. (c) Distribution of the number of differential interactions across all chromosomes. (d) Chromosomes 14, X, and 7 exhibited the greatest interaction changes with most rearranged regions indicated. (e) A ranking of gene promoters with most differential interactions on Chromosome 7, in which Tgfb1 was ranked near the top (see insert). (f) Interaction landscape at the Tgfb1 region on Chromosome 7 revealed interweaving patterns of interaction gains (green) and losses (red). Certain sites harbored more interaction changes than others. Notable was Tgfb1-Ltbp4 loop rearrangement (circled) which also overlapped with CTCF, MyoD, ATACseq peaks. Rearrangement was formed between Tgfb1 (highlighted by yellow line), and Ltbp4 (yellow line) and bore the signature of adjacent gains and losses at the same location.

To evaluate the effects of Matr3 depletion on chromatin architecture at steady-state, we performed Hi-C using Matr3 KO bulk on Day 0. Indeed, rearranged loops were observed upon Matr3 loss (Supplemental Figure 6). Moreover, changes in chromatin loops were correlated with differential open chromatin accessibility (Supplemental Figure 7). Taken together, these results provide strong evidence that Matr3 is essential in maintenance of chromatin loops around chromatin accessible regions of the genome.

### Loss of Matr3 disrupts YY1 binding and YY1-enriched cohesin loading

Chromatin loops are comprised of structural chromatin loops, and, on a finer scale, long-range chromatin interactions, including enhancer-enhancer (E-E), enhancer-promoter (E-P) and promoter-promoter (P-P) loops (Rao et al., 2014, Schoenfelder et al., 2015, Hsieh et al., 2020). The strength of E-P and P-P interactions positively correlates with the level of gene expression (Hsieh et al., 2022). Yin Yang 1, YY1, a ubiquitously expressed transcription factor that preferentially occupies enhancers and promoters in mammalian cells, is a critical regulator of E-P loops, whereas CTCF is enriched at insulator elements (Weintraub et al., 2017, Wang et al., 2018). Matr3 was found in complexes with CTCF and cohesin; moreover, CTCF and cohesin occupancy was decreased upon Matr3 loss (Cha et al., 2021). Given the reported association between YY1 and cohesin (Weintraub et al., 2017, Hsieh et al., 2022), we hypothesized that Matr3 might impact chromosomal loops by affecting cohesin and YY1 occupancy. By CUT&RUN analysis (4hrs. dTAG47 treatment), we observed that YY1 binding was enhanced at 2840 locations (compared to 947 locations where YY1 was decreased) upon Matr3 depletion (Figure 5a). To interrogate potential interactions between YY1 and cohesin Rad21, we mapped the pattern of redistribution of Rad21 binding after Matr3 depletion (4hrs. dTAG47). Acute removal of Matr3 was associated with reduced Rad21 occupancy adjacent to enhanced YY1 binding sites (Figure 5b, c). This pattern of enhanced YY1 binding and loss of Rad21 binding was observed at 208 genomic loci, corresponding to roughly 19% of sites at which Rad21 binding was lost (1127 sites). These observations suggest that loss of cohesin Rad21 following depletion of Matr3 might be compensated in part by increased YY1 occupancy. In addition to decreased Rad21 occupancy, sites of differential YY1 binding were often found in association with increased MyoD occupancy (Figure 5d) and loss of chromatin accessibility (Figure 5e). We suggest, therefore, that Matr3 loss perturbs connections between the transcriptional machinery and the higher order chromatin structure in part through changes in YY1 binding.

**Figure 5.**
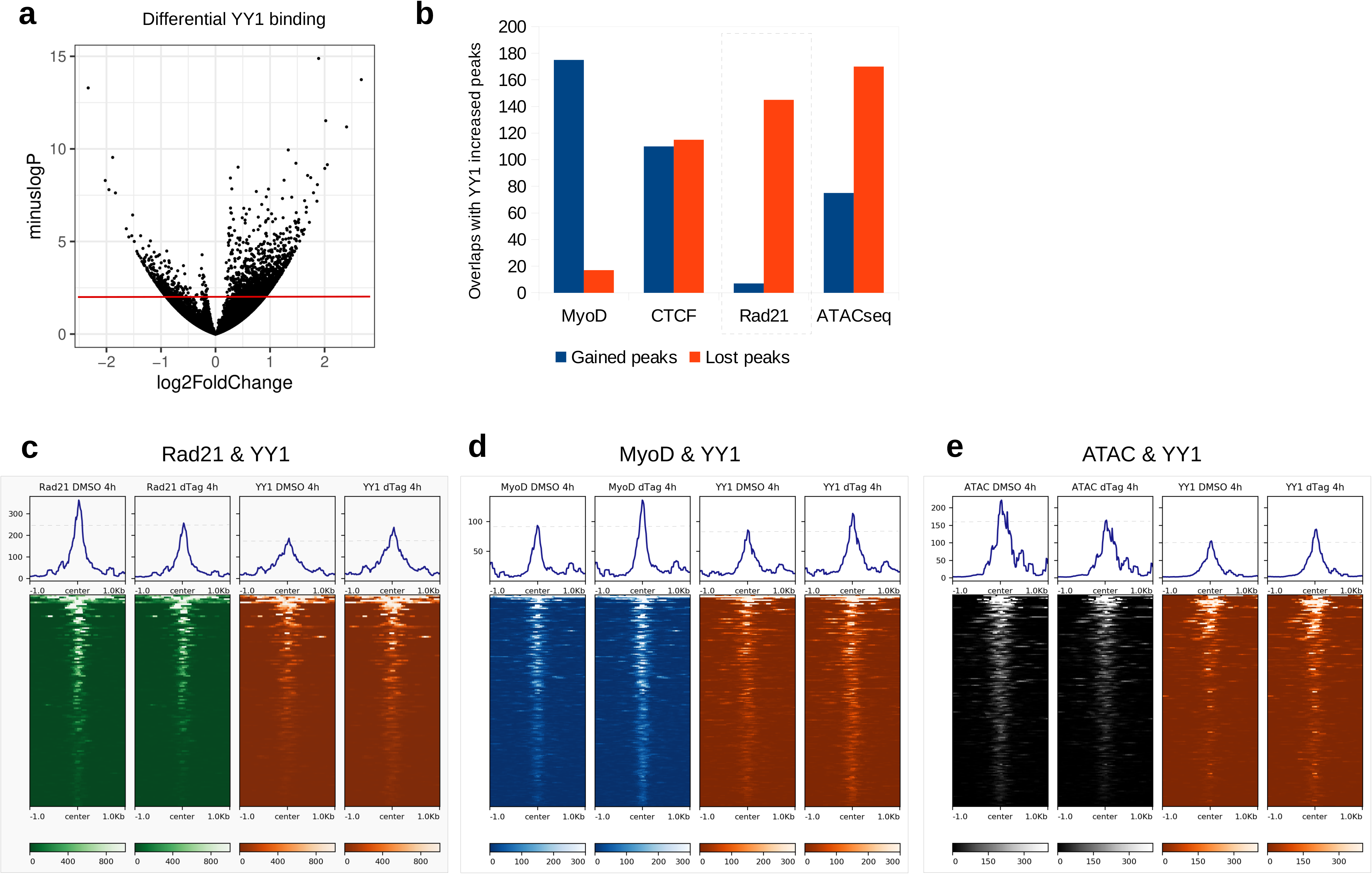
Loss of Matr3 disrupts YY1 binding and YY1-enriched cohesin loading. (a) YY1 binding enhanced upon Matr3 depletion (4hrs.). Volcano plot showing the logFoldChange vs. -logPvalue of YY1 binding changes (measured by Δ=dTAG-DMSO treatment). Red line: significance line (P=0.05). (b) Co-occurrence of differential YY1 occupancy with MyoD, CTCF, Rad21 bindings, ATAC-seq. Number of shared differential peaks between YY1 increased (4hrs.) and the gained/lost portion (4hrs.) of MyoD, CTCF, Rad21, or ATAC-seq. Box: the overlap percentage with Rad21 binding loss was highest, showing a relationship between YY1 differential binding and cohesin loss. (c) Heatmap showing that Rad21 binding was lost at sites of YY1 binding increase. Dashed line on the histogram shows the reference line to aid comparison between dTAG47 treatment and DMSO. (d) Heatmap showing MyoD binding was enhanced at sites of YY1 binding increase. (e) Heatmap showing ATAC-seq peaks were decreased at sites of YY1 binding increase.

### Loss of Matr3 disrupts YY1-mediated enhancer-promoter loop formation

As YY1 is a primary regulator of EP loops (Weintraub et al., 2017), we hypothesized that loss of Matr3 would elicit changes in the chromatin loop domains. We next investigated YY1-mediated chromatin loop changes upon depletion of Matr3 (4hrs., dTAG47). Loci characterized by enhanced YY1 occupancy exhibited a greater number of increased E-E and E-P loops. The Mphosph8 locus illustrates these changes (Figure 6a). Increased interactions were observed between segment 1 and segments 2, 3, 4, while decreased interactions were detected only at segment 5. In the entire 1.5Mb region, we detected 11 increased interactions as compared with 4 interactions that were decreased.

**Figure 6.**
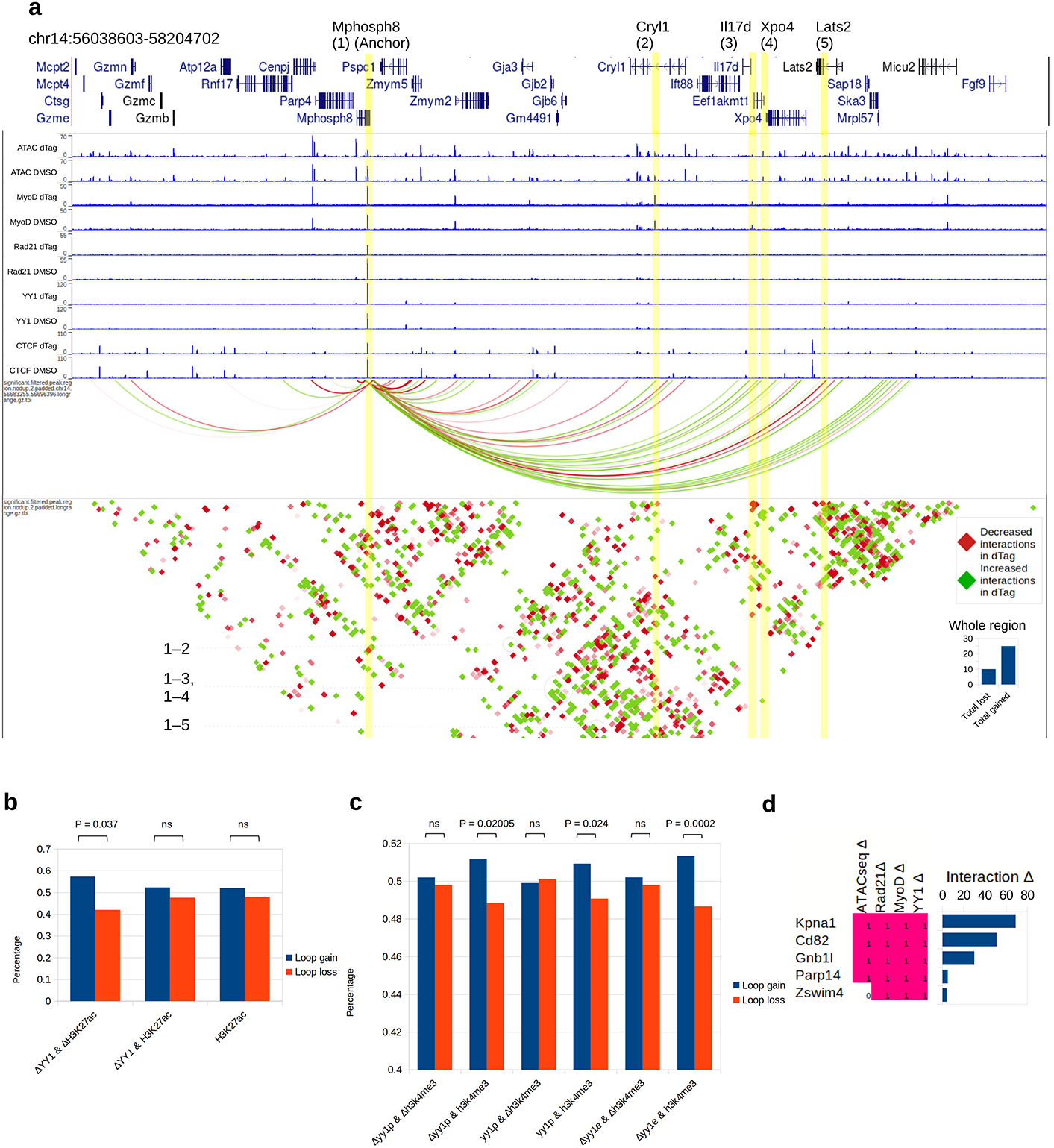
Loss of Matr3 disrupts YY1-mediated enhancer-promoter loop formation. (a) Loci characterized by enhanced YY1 occupancy exhibited increased E-E and E-P loops. Mphosph8 locus was an example of a promoter with differential YY1 binding causing EP-loop gains with neighboring MyoD, ATAC-seq peaks. Mphosph8 (highlighted by yellow line) exhibited a dual YY1-gain and Rad21-loss signature. The gene formed increased interactions (highlighted by green arcs) with Cryl1 (2), Il17d (3), Xpo4 (4), and Lats (2) (highlighted by yellow lines), which displayed MyoD and ATAC peaks at these positions. Interaction dot map further reaffirmed the positive increase in EP-loop interactions. Histograms (bottom right) indicated the interaction gain/loss scores across the whole region in the plot. (b) YY1-mediated EP loops changed in a H3K27ac-dependent manner in the absence of Matr3. Loop gain/loss ratios were measured. YY1 enhancers with joint H3K27Ac change (ΔYY1 & ΔH3K27Ac) were more likely to demonstrate loop gains than YY1 enhancers with no H3K27Ac changes (ΔYY1 & H3K27Ac). (c) Increased YY1 and differential H3K4me3 sites were associated with reduction of E-P loop gain. YY1-mediated EP loops changed in a H3K4me3-dependent manner in the absence of Matr3. YY1p: YY1 promoters. YY1e: YY1 enhancers. YY1 promoters with co-differential H3K4me3 (ΔYY1p & ΔH3K4me3) were compared to YY1 promoters with no H3K4me3 changes (ΔYY1p & H3K4me3). (d) Sites of 4-way (ATAC-seq, Rad21, MyoD, YY1) co-differential signatures were more likely to harbor loop interaction changes.

Genome-wide, increased looping upon Matr3 depletion was observed more frequently where increased YY1 occupancy was detected at the enhancer (E) anchor of differential E-P loops, marked by H3K27Ac. Furthermore, enhancers marked by increased YY1 and altered H3K27Ac deposition exhibited a higher percentage of gained interactions than lost interactions (the gain/loss ratio was 196:146, bias: 1.36, see Figure 6b, ΔYY1&ΔH3K27Ac). This bias towards gained interactions was much stronger than gained YY1 sites with unaltered H3K27Ac (the gain/loss ratio was 1368:1246, bias: 1.09, see ΔYY1&H3K27Ac), or all H3K27Ac sites (the gain/loss ratio was 11312:10430, bias: 1.06, see H3K27Ac). Thus, Matr3 depletion is associated with gained E-P loops at enhancer YY1 binding sites marked by H3K27Ac deposition.

An alternative scenario was observed in which E-P loops coincided with changes in active marks H3K4me3 at gene promoters. We analyzed the E-P loops of this category of genes, where H3K4me3 was gained at the gene promoter and enhancers were occupied by YY1 (Figure 6c). We observed more balanced E-P loop changes radiating from such promoters (gain/loss ratio is 1.01) (see Δyy1e&Δh3k4me3 compared with Δyy1e&h3k4me3 in Figure 6c). This reduction of E-P gains at differential H3K4me3 sites may reflect greater direct promoter-based regulation than E-P loop-based gene regulation. Taken together, these data reflect diverse H3K27Ac-based and H3K4me3-based mechanisms by which Matr3 depletion elicited changes in the YY1-mediated E-P loop landscape.

Given that changes in MyoD and Rad21 occupancy, and in chromatin accessibility, as assessed by ATACseq, appeared to be strongly associated with YY1, we hypothesized that there might be a subset of binding sites with co-occurring changes most associated with such interactions. We computed genome-wide the number of loci characterized by 4-way differential chromatin accessibility and MyoD, Rad21, and YY1 binding, and the number of interaction differences at these locations, as revealed by Hi-C. Overall, we identified 17 loci that have interaction differences. 3 loci were prioritized on the basis of most altered interactions (Figure 6d). Among the candidates, Cd82 and Kpna1 were notable for their involvement in muscle stem cell activation and proliferation with links to skeletal muscle defects (Hall, Arielle et al., 2020, Choo, Hyo-Jung et al., 2016).

Depletion of Kpna1 was associated with premature activation of muscle stem cells, and subsequent proliferation, apoptosis, and satellite cell exhaustion (Choo, Hyo-Jung et al., 2016). Indeed, the top 3 genes (Kpna1, Cd82, Gnb1l) (Figure 6d) that exhibit 4-way co-differential marks along with strong loop interaction changes all displayed differential gene expression (P=0.0531, P=0.054, P=0.0029 respectively, Day 0).

Taken together, we propose a model to account for how loss of Matr3 affects gene expression and differentiation (Figure 7). We posit that Matr3 loss destabilizes cohesin-CTCF complex binding by reducing cohesin loading, therefore altering chromatin structural loops and affecting long distance interactions. As a mechanism for partial compensation of cohesin loss, YY1 is recruited to chromatin sites, thereby disrupting the E-P loop landscape. During myogenesis, gain of YY1-mediated E-P loops may contribute to recruitment of MyoD, leading to gene expression changes as skeletal muscle development progresses.

**Figure 7.**
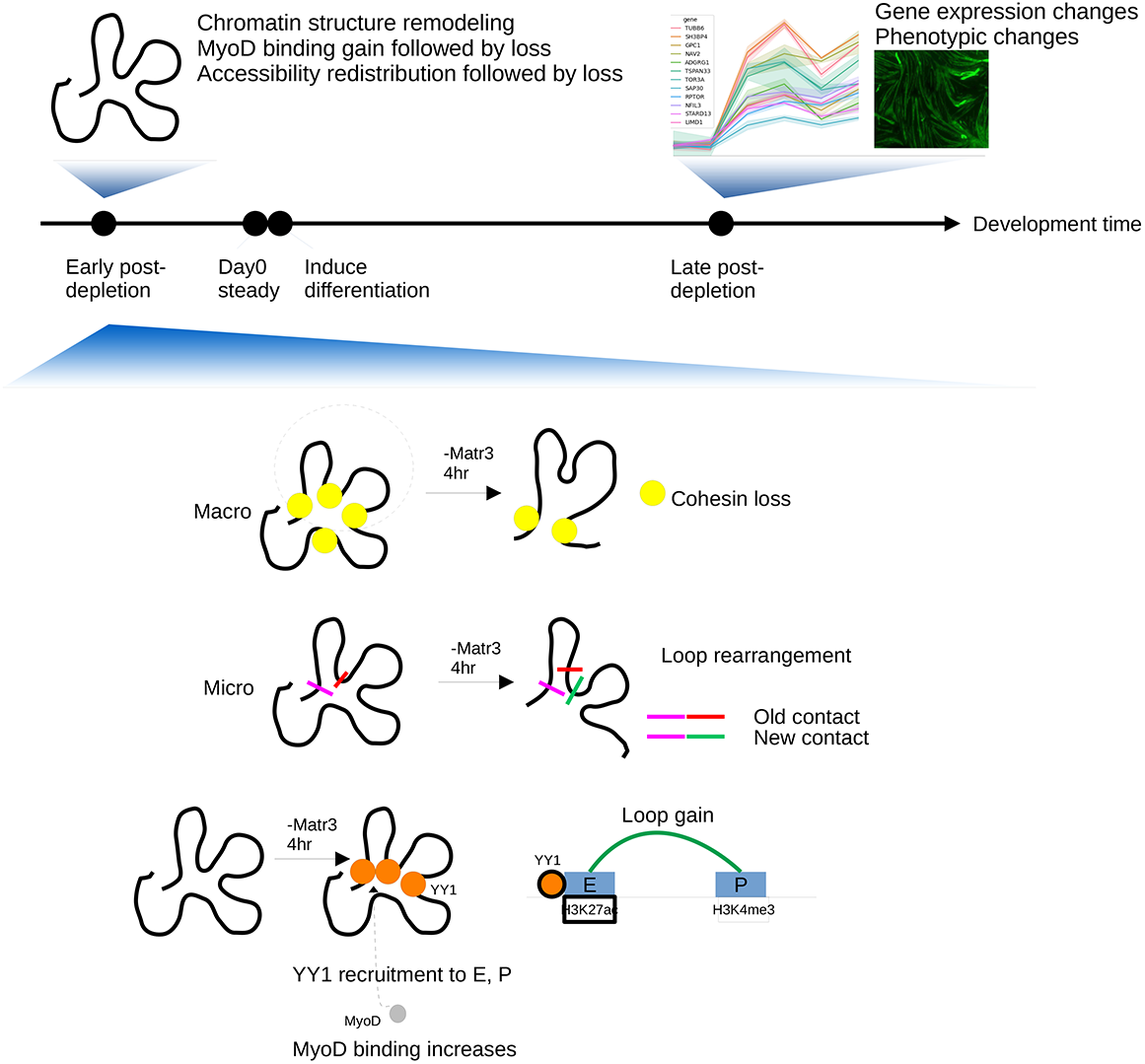
Matr3 mediates differentiation through stabilizing chromatin accessibility and chromatin loop-domain interactions, and YY1 mediated enhancer-promoter interactions. Acute depletion of Matr3 loss immediately reduced cohesin loading, and destabilized chromatin structural loops and affected long distance interactions. As a mechanism for partial compensation of loop re-arrangement, YY1 was recruited to sites of cohesin loss, as well as enhancers and promoter sites, and formed new E-P loop, thereby disrupting the E-P loop landscape. During myogenesis, MyoD was recruited to those chromatin sites, leading to gene expression changes as skeletal muscle development progressed.

## Discussion

Matr3 has been traditionally ascribed with roles in RNA splicing and transport (Coelho et al., 2015, Salton et al., 2011). Our prior work demonstrates that Matr3 also associates with chromatin through interactions with architectural proteins CTCF and cohesin/Rad21, and orchestrates developmental transitions in embryonic stem cells and erythroid cells (Cha et al., 2021). Here, through studies of Matr3 contributions in the context of myogenesis, we extend a conserved role for Matr3 in differentiation and explore in detail how its loss impacts chromosome looping.

With the goal of probing the direct roles of Matr3 in chromatin structure, rather than secondary effects resulting from its absence, we established an experimental platform for acute targeted protein degradation (TPD) of Matr3 in myogenic C2C12 cells by introducing a modified FKBP cassette at high efficiency into the Matr3 locus. Following addition of the PROTAC dTAG47, Matr3 was depleted within 4 hrs. in bulk cell populations. This strategy permitted direct assessment of the consequences of Matr3 loss on dynamic aspects of chromatin organization. Several new perspectives on the role of Matr3 arose from these studies.

First, upon TPD of Matr3 in C2C12 cells, we uncovered dynamic, early changes in overall chromatin accessibility and MyoD occupancy. Specifically, we observed mixed early gains in chromatin accessibility and predominant gains in MyoD occupancy, which were followed by loss of both accessibility and MyoD occupancy at later times (Figure 3a-b). These early changes in chromatin accessibility and MyoD occupancy were not reflected in rapid changes in gene expression, but were associated with changes later in muscle differentiation (Figure 3h-i).

Second, our findings revealed that gene expression changes are manifested days later than the initial perturbation of chromatin structure elicited by TPD of Matr3. This observation is in accord with a recent study, which demonstrated that an RNA-binding protein complex containing MATR3 assembled at an inactive X (Xi)-compartment mediated gene repression even after day 3 of differentiation (Pandya-Jones et al., 2021), at which point the Xist RNA had disappeared. Thus, a seeded protein condensate may have long lasting and sustained effects on gene expression (Pandya-Jones et al., 2021). In our study, we observed that genes co-expressed and found in a Matr3-complex were cumulatively perturbed upon TPD of Matr3 and subsequent C2C12 cell differentiation (Supplemental Figure 4). These findings highlight how direct effects on chromatin architecture may only be appreciated at later times in cell differentiation.

Third, chromosomal loop rearrangements observed following TPD of Matr3 were broad in scale, reflecting a general rather than locus-specific role of Matr3 in maintaining chromatin architecture. Rearrangements included simultaneous gains of new enhancer-promoter, enhancer-enhancer loops, and losses of adjacent existing loops (Figure 4b). We hypothesized that loop rearrangements upon Matr3 loop are facilitated in part by the sliding of cohesin rings and YY1 occupancy near enhancer and promoter sites. We observed that acute Matr3-depletion preferentially affected sites at which both cohesin and YY1 were bound (Figure 5b-c), and altered YY1-mediated EP loops in a histone H3K27Ac and H3K4me3-dependent manner (Figure 6b-c). Sites of differential H3K27Ac and H3K4me3 marking harbored more EP-loop changes. Previously, YY1 was reported to directly regulate enhancer-promoter interactions (Weintraub et al., 2017). Our data provide evidence that regions of YY1-binding perturbed upon Matr3 loss exhibit E-P interaction differences. Given the observed overlap of differential MyoD/Rad21/YY1/accessibility sites, we speculate that upon rapid TPD of Matr3, YY1 may be recruited to sites with gained chromatin accessibility/MyoD occupancy, and differential histone acetylation to form EP loops and activate downstream gene targets (Figure 7).

Matr3 has been reported to interact with members of the spliceosome complex and serve a role in RNA splicing (Salton et al., 2010). Perturbation of the spliceosome member Hnrnpm led to reduced Matr3, and knockdown of AKAP8, a member of the Hnrnpm RNA splicing complex, was associated with a mesenchymal phenotype and accelerated EMT (Ramesh et al., 2020, Hu et al., 2020). These findings indicate that perturbing a member of the RNA splicing complex may have developmental consequences at later times. Taken together with our observations of the role of Matr3 in maintenance of chromatin architecture, it would be interesting to explore whether depletion of other members of the spliceosome complex elicit changes in chromatin structure and accessibility.

Mutations and dysregulation of Matr3 have been associated with ALS and Spinal Muscular Atrophy (SMA) (Chi et al., 2018). While it has been generally accepted that these effects reflect perturbation of processes related to RNA processing/splicing (Xiao et al., 2012, Singh et al., 2018), our findings implicate chromatin loop level changes as an alternative, or contributing mechanism. For example, upon Matr3 depletion we observed rearrangement within the Ltbp4-Tgfb1 locus (Figure 4f), a region linked to muscular dystrophy in GWAS (Heydemann et al., 2009, Lamar et al., 2016, Demonbreun et al., 2021). Future efforts may be directed to investigate the significance of regulatory elements involved in long-range chromatin interactions which are perturbed upon Matr3 loss. The list of chromatin loop changes prioritized by Hi-C upon TPD of Matr3 may be used to reinterpret susceptibility loci from neuromuscular disease GWAS studies.

In summary, our work reveals a critical role of Matr3 in development in several contexts, and directly links the alteration in proteins within nuclear compartments to changes in chromosomal architecture. Through effects on chromatin accessibility, and the occupancy of master transcriptional regulators and YY1, Matr3 impacts chromatin structure to orchestrate differentiation.

## ACKNOWLEDGMENTS

We thank the Molecular Biology Core Facility at DFCI, HESC Core Facility, BCH/PCMM Microscopy Core, and Viral core at BCH. We thank Drs. Nan Liu for CUT&RUN reagents and discussion, Manuela Gussoni for myotube immunostaining antibodies and discussion, Louis M. Kunkel Lab for DMD antibody, Stephen Tapscott Lab for MyoD antibody, Kevin P. Campbell and Mary E. Anderson for DMD complex antibodies and analysis, and Jun Qi for dTAG47. Computational analyses were conducted on the O2 High Performance Compute Cluster, supported by the Research Computing Group at Harvard Medical School. Stuart H. Orkin is an Investigator of the HHMI.

## AUTHOR CONTRIBUTIONS

T.L, Q.Z. and S.H.O. designed the study. T.L. performed the experiments, and analyzed and interpreted the data. Q.Z. performed computational analyses and interpreted the results. K.Y. analyzed Hi-C compartments. G.Y. helped with computational analyses tools. T.B. performed Yokogawa CV7000 imaging analysis. T.M.S. helped Yokogawa CV7000 imaging analysis tools. H.C. generated the plasmid for Matr3 knockout clones. S.M. helped design the plasmids for knock-in dTAG-Matr3.

S.H.O. supervised the work. T.L., Q.Z. and S.H.O. wrote the manuscript, with input from all the authors.

---

**Supplemental Figure 1 Generate *Matr3* KO C2C12 using CRISPR/Cas9.** (a) A schematic depiction of using CRISPR/Cas9 to delete the entire Matr3 gene body to generate Matr3 KO clones (refer to method section). bi-allelic Matr3 deletion clones are screened by PCR. (b) Matr3 KO clones were confirmed by western blots. 2 independent Matr3-KO clones were used. (c) Matr3 KO clones were confirmed by immunostaining. C2C12 cells (Day0) were immunostained with DAPI (blue) and Matr3 (red). scale bar 20um. (d) Clonal variation in growth and differentiation among C2C12 clones. 4 representative wildtype clones and 2 Matr3-KO clones were differentiated and monitored for 4 days.

**Supplemental Figure 2. Matr3 KO bulk was generated using a high-efficient guide.** Synthego ICE Sequencing report showing Matr3 KO bulk using single guide delivered as RNP had a high indel and knockout score.

**Supplemental Figure 3. N-terminal dTAG-Matr3 bulk was generated and confirmed.** (a) Biallelic dTAG-Matr3 knock-in clones were screened by PCR. >85% of the clones from the screening were bi-allelic dTAG-Matr3 knock-in. (b) High-efficient knock-in bulk FKBP F36v-Matr3 were sorted by GFP and confirmed by western blots. Cells enriched in the top 0.5% highest GFP signal were sorted (GFP-sorted). FKBP F36v-Matr3 (refer as dTAG-Matr3) bands were enriched in the GFP-sorted bulk (left lane), compared with unsorted bulk (middle lane), indicating the high-efficient knock-in efficiency. (c) dTAG-Matr3 protein was degraded by all 3 PROTACs (dTAG-v-1, dTAG13, dTAG47 500nM) completely after 24hrs. treatment. (d) dTAG-Matr3 in C2C12 were degraded more efficiently by dTAG47 (500nM, 4h) than dTAG-v1 (500nM, 12h).

**Supplemental Figure 4. Matr3 co-expressed genes are perturbed in expression at later time points**. First, we used SEEK to identify Matr3-coordinated (or co-expressed genes). We entered MATR3 along with 4 other genes found in co-complex with MATR3 (TARDBP, MATR3, CELF1, PTBP1) forming a coherent query gene-set, in order to steer SEEK to use the right datasets (where MATR3 is active) for co-expression analysis. SEEK returned the co-expressed genes to the query using a large body of gene expression data sets where MATR3 and complex expression was high. The top 500 co-expressed genes were isolated and correlated with our C2C12 Day 0, Day 4, Day 6 differentiated KO vs. WT gene-sets, in order to assess what proportion of the co-expressed genes changed expression during development. As shown, Matr3 co-expressed genes were increasingly perturbed in expression over development time.

**Supplemental Figure 5. Matr3 has minimal impact on compartments switching and interaction.** (a) Heatmap showing substantial reduction of Rad21 binding in Matr3 KO. CTCF occupancy was marked by both gains and losses in Matr3 KO. (b) Snapshot of compartmentalization landscapes of indicated samples at Chromosome 1. PC1 values of Hi-C data were shown, with red color representing active compartment A and blue representing inactive compartment. (c) Saddle plots showing the average interaction within and between compartments A and B. Numbers in the heatmaps indicated the average compartment strength (Log2 Obs/Exp) for intra- (A-A in top-left, B-B in bottom-right) and inter-compartment interactions (AB, top-right or bottom-left). The differential heatmap for each comparison was shown on the right panel with red/blue color representing increase/decrease in interaction strength.

**Supplemental Figure 6. Re-arranged loops were observed in the WT and KO Matr3 Hi-C experiments.** Gene loci on chromosome 7 were arranged based on differential loop interaction scores as observed in the WT vs. KO Matr3 Hi-C experiments. **Top**: lost loop interaction scores. **Bottom**: gained interaction scores at each gene locus. Loop interaction scores were summarized per gene TSS location. **Boxed**: Tgfb1 locus was again ranked among top of the most rearranged gene loci.

**Supplemental Figure 7. Differential ATAC-seq signals are correlated with differential Hi-C loop interactions.** (a) We first identified all Hi-C anchors that overlap with ATAC-seq peaks. Then, we interrogated the strength of interactions between Hi-C anchors that overlap with ATAC-seq peaks. For a pair of interacting loci (x, y), where x is anchor 1 and y is anchor 2, both of which overlapping with ATAC-seq, the observed interactions for (x, y) in KO was subtracted from the observed interactions of (x, y) in WT, and the difference was plotted (green shade: positive, red shade: negative). Thus, this analysis correlated with differential Hi-C interaction signals. Results were sorted based on WT/KO ATAC-differential P-values of individual anchors. Results were sorted into 25 bins, each bin stores 5% of total ATAC-seq peaks. (b) The top 10% fraction, divided into 10 1% bins. Each histogram quantified the number of differential Hi-C interactions observed in that percentile bin.

**Supplemental Figure 8. Loop calling with HIFI pre-processing.** (**a**) Loop calling at various resolutions and various levels of stringencies using Hiccup algorithm, with HIFI-based preprocessing. 5kb, 10kb, 25kb loops and 50kb loops were visualized within 10 kb-, 20 kb-, 50 kb- and 100 kb-sized windows respectively. Three stringencies of loops (1.0, 1.2, and 1.5 as determined by Hiccup score) were selected for visualization. (**b**) Loop calling comparison, showing agreement between Hiccup and FitHiC with and without HIFI imputation. (**b1**) With HIFI imputation, the overlap (by Jaccard coefficient) between Hiccup and FitHiC detected loops increased from 6% to 13% (for 10kb loops), 17% to 30% (for 25kb loops), and 27% to 35% (for 50kb loops). (**b2**) With HIFI imputation, the number of agreed, detected loops per 10Mb also increased from 30 to 200 (for 5kb), 110 to 270 (for 10kb), 120 to 400 (for 25kb), and 80 to 350 (for 50kb), representing on average 3-5 fold increase.

## Method and Materials

### C2C12 Cell culturing and differentiation assay

The C2C12 cell line was obtained from the American Type Culture Collection (ATCC® CRL1772™). C2C12 cells were grown and differentiated following the standard protocol (Doynova et al., 2017). C2C12 myoblasts were propagated at 37°C, 5% CO_2_ in Dulbecco’s modified Eagle’s medium (Thermo Fisher Scientific) supplemented with 10% fetal bovine serum (Omega Scientific) and antibiotics (penicillin 100 U/ml, streptomycin 100μg/ml, Thermo Fisher Scientific).

For differentiation, high-density cultures (∼80% confluency) were switched to differentiation media (DMEM, 2% horse serum (HS, Gibco®16050–122), penicillin 100 U/ml, and streptomycin 100μg/ml) to induce myotube formation. Myotubes at different times (0, 2, 4, 6 days) post differentiation were used for further analysis.

### Generation Matr3 KO clones by CRISPR/Cas9

CRISPR/Cas9 editing was used to delete the entire Matr3 gene body in C2C12 cells (Cha et al., 2021, Bauer et al., 2015). Briefly, paired single guide RNAs (sgRNAs) for site-specific cleavage of genomic regions were designed upstream and downstream of Matr3 coding region. Annealed oligos were ligated into the pX330 vector using a Golden Gate assembly strategy. A pair of CRISPR/Cas9 constructs (5 μg each) targeting each flanking region of the deletion site was transfected into 5×10^5^ cells with 0.5 μg of GFP expression plasmid using BTX Harvard Apparatus. The top ∼0.2% of GFP positive cells were sorted by FACS 48 hours post-transfection. Single cell derived clones were isolated and screened for biallelic deletion of target genomic sequences.

The gRNA sequences and the primers used for genotyping the clones were listed in Supplemental Table of primers and antibodies.

### Generation Matr3 KO bulk using Cas9/RNP transfection

RNP electroporation with 4D-Nucleofector™ were performed according to 4D-Nucleofector™ Protocol C2C12 cells (Lonza). Freshly thawed C2C12 cells were recovered and subculture 2 days before Nucleofection. Cells were harvested by trypsinization (0.25% trypsin, 37°C) and neutralization with culture medium. Next, cells were washed once with PBS and separated to 2 x 10^5^/sample. Cas9/sgRNA RNP was generated by mixing 120 pM guide RNA (20-O-methyl analog and 30 phosphorothioate internucleotide modified-sgRNA synthetic from Synthego, with 61 pM Cas9-Alt-R protein (IDT), and incubated at room temperature for 15min. Meanwhile, 2 x 10^5^ cells/sample were collected by centrifugation at 90xg for 10min, and resuspended carefully in 20ul 4D-Nucleofector™ Solution (SE Cell Line 4D-Nucleofector™ X Kit, Lonza). Thereafter, RNP mix was added to the cells, and nucleofection was carried out in a 4D-nucleofector X Unit with program CD-137. Immediately after nucleofection, cells were supplemented with 80 ul prewarmed culture medium. Cells were then split to two wells of 6-well plate filled with 1.5ml culture medium. Knockout efficiency using PCR and western blots were performed 2 days post RNP delivery. The gRNA sequences and the primers used for genotyping the clones are listed in Supplemental Table of primers and antibodies.

### Generation knock-in N-terminal dTAG-Matr3

#### Plasmid design

N-terminal fusions of FKBP^f36v^ to endogenous matr3 were generated with double-strand break using Cas9/sgRNA followed by delivery of HDR donor cassette via crude-rAAV preparations as described elsewhere (Mehta et al., 2023). At the N-terminus, eGFP-p2a-FKBP^f36v^ was inserted after the ATG start codon. In instances where the cut site of a gRNA was distal to the intended insertion site, the wobble bases of all amino acid sequences were re-coded to prevent unintended repair. Homology arms were designed to be ∼900bp for the N insertions.

### Generation of crude rAAV preps

Crude rAAV viral preps were generated as described (Mehta et al., 2023). Briefly, HDR templates consisting of eGFP-p2a-FKBP^f36v^ flanked by homology arms were chemically synthesized ordered and cloned between the two Inverted Terminal Repeat (ITR) sequences of a pAAV2 vector to generate a transfer vector. Thereafter, the triple-transfection method was used to generate crude rAAV lysates (Robert et al. 2017).

### Generation dTAG knock-in C2C12 clones

C2C12 cells were electroporated with Cas9/sgRNA complexes targeting the HDR insertion site (with sgRNA protospacer sequence spanning the insertion site). Electroporation was performed using the same setting as the Cas9/RNP transfection to generate Matr3 KO bulk, following the protocol of 4D-Nucleofector™ Solution (SE Cell Line 4D-Nucleofector™ X Kit, Lonza). In this setup, Cas9/sgRNA Ribonucleoprotein (RNP) mixture was generated with 120 pM guide RNA (2’-O-methyl analog and 3’ phosphorothioate internucleotide modified-sgRNA, custom ordered from Synthego Corp., with 61 pM Cas9-Alt-R protein (IDT). Immediately after RNP electroporation, 150ul crude rAAV viral prep (see above) was added to the cells (∼10% of total culture medium), and the cells were returned to the 37°C incubator for 3 days before expansion.

One week after electroporation/ AAV transduction, cells were sorted and top 0.2% GFP of parent cells were collected to generate the dTAG-Matr3 knock-in bulk and screen for single clones. To isolate single-cell clones, ∼30 cells were seeded per 96-well plate. Single clones were expanded and genotyped for the desired knock-in by PCR on genomic DNA extracted with Quickextract solution (Lucigen). Clones showing biallelic knock-ins were expanded and frozen in LN2 at early passages. The gRNA sequences and the primers used for genotyping the clones are listed in Supplemental Table of primers and antibodies. Clones and bulk were validated by Western blot and Sanger sequencing.

### Western blot

Western blots were performed as described in previous paper with modification (Liu, et al., 2017, Bi et al., 2017). Protein was isolated from cells or myotubes using RIPA buffer (Boston Bioproduct BP115 containing 50 mM Tris, 150 mM NaCl, 10% glycerol, 1% NP40, 0.5% sodium deoxycholate, 0.1% SDS) with 0.5 mM PMSF (phenylmethylsulfonyl fluoride), 1 mM Na_3_VO_4_, and 1 mM NaF and complete EDTA-free protease inhibitor mixture (Roche Applied Science). Cell lysates were disrupted mechanically by passing them through 25G 5/8 needles 10 times and then centrifuged at 10,000g for 5min. The supernatant was collected, and protein concentrations were determined with BCA protein Assay Reagents (ThermoFisher Scientific, 23225) followed by measurement with NanoDrop. Protein samples were mixed with 4X Laemmli sample buffer (BIO-RAD # 161-0747) and same amount of protein (10ug-40ug) was loaded and separated by Mini-PROTEAN® TGX™Precast Gels (4%-20%), Blots were probed with primary antibody and the Horseradish peroxidase conjugated secondary antibodies (antibodies used in western blots could be found in supplemental table of primers and antibodies). Immunoblots were visualized using ECL plus reagent (GE Life Sciences), and the Fiji program was used to quantify protein abundance.

### Immunohistochemistry

For immunofluorescence staining with C2C12 myoblast and myotubes, 4-well Permanox Chamber Slides (Nunc™ Lab-Tek™ Chamber Slide System 177437, Thermo Fisher) were used for Olympus confocal imaging, and Greiner CELLSTAR® 96 well plates (Greiner 655090, Sigma) were used for Yokogawa CV7000 microscope. Cells were seeded on the plates/slides that were pre-treated with 0.3% gelatin, and kept in incubator for proliferation and differentiation. Cells were rinsed with PBS three times and then fixed with 4% PFA in PBS for 15 min at room temperature. After fixation, cells were rinsed twice with PBS before permeabilization with 0.25% Triton X-100 in PBS for 10 min. Next, cells were blocked with freshly made 5% normal donkey serum (or 5% goat serum) in PBST (PBS with 0.25% Triton X-100) for 1 h at room temperature, and were incubated with primary antibody in blocking solution overnight at 4 °C. The next day, after three washes with PBS, cells were incubated with secondary antibody in PBS for 1 h in dark at room temperature. After three washes with PBS, the cells were mounted in VECTASHIELD® Antifade Mounting Medium with DAPI (H-1200-10).

### Imaging and image analysis

#### Olympus FV1000 confocal microscope imaging and analysis

To get higher resolution of cell nucleus, cells were seeded on 4-well Permanox Chamber Slides (Nunc™ Lab-Tek™ Chamber Slide System 177437) and imaged with an Olympus FV1000 confocal microscope as described previously (Liu et al., 2017). Confocal stacks were imaged with Olympus FV1000 equipped with 20×/0.85 N.A. and 100× 1.40 N.A. oil-immersion objectives. For double-labeling experiments, sequential scans of argon ion 488-nm and HeNe (633 nm for AlexaFluor 647) lasers were used to avoid bleed through between channels. The Fluoview “Hi-Lo” look-up table was used to set the maximal signal below saturation and set the background to near zero using the high voltage and offset controls. Z-series images were obtained at a 2-μm step size, and Kalman averaging was not used. Original images were saved as 12-bit OIB format and processed using FV1000 confocal software to generate maximum intensity projections (Z-projections). Images were adjusted for brightness and contrast using ImageJ/Fiji software. For each genotype, images were acquired using the same settings (power, gain, offset) at the same time.

### Yokogawa CV7000 microscope imaging and analysis

For high-throughput analysis, imaging was performed on a Yokogawa CV7000 microscope with 10x imaging using appropriate filter and laser settings (DCAF8/DMD -AF488: 488 excitations with 510/20 BP filter, MHC - AF647: 640 excitations with 661/20 BP filter). Images were acquired with an exposure of 1000ms for all channels and captured with 2x2 binning at +/-20um from the focal point with a step size of 10um (determined imaging parameters required to capture the whole depth of the muscle fiber). These images were automatically summed stacks in the CV7000 software and exported for analysis in Fiji, and subsequently data interpretation in Python (v3.85).

In order to create a per-channel light path correction image, multiple empty wells were stained and treated with anti-fade and were median projected. The per-channel light path correction image was subtracted from the respective channel in all experimental wells. Subsequently, all AF647 and AF488 images were added together in order to discover all forming muscle fibers within the well. A gaussian blur was applied to this image, and maxima were identified. These maxima were then increased in size in order to establish ROIs within the muscle fibers. The average per-channel fluorescence intensity was then recorded per ROI and imported to Python for analysis. When plotting MHC levels, it was discovered that the distribution was multimodal with a large peak in the intermediate MHC expressing sub-population. High MHC levels (>10,000 Fluorescence intensity) tended to arise from muscle fibers that were folded or overlapped, while extremely low levels (<1000 Fluorescence Intensity) were usually a result of improper muscle fiber identification. Therefore, in order to establish higher levels of consistency, we chose to focus on ROIs within the intermediate MHC distribution (an MHC Fluorescence intensity between 1,000-10,000). Note that there was not a significant difference in the distribution of MHC within ROIs when compared between WT and Matr3 KO. After gating this data, the corresponding bar plots were generated using the Seaborn. Analysis of statistical significance was performed using scipy Welch’s t-test and Mann-Whitney U test with a p-value of .05 and assumption that KO fluorescence of the Target was less than the WT fluorescence of the Target, where the target is either DMD or DCAF8.

### RNA-seq

Total RNA was isolated from cells using the RNeasy Plus Mini Kit (Qiagen). RNA sequencing libraries were prepared using Roche Kapa mRNA HyperPrep sample preparation kits from 100ng of purified total RNA according to the manufacturer’s protocol. The finished dsDNA libraries were quantified by Qubit fluorometer, Agilent TapeStation 2200, and RT-qPCR using the Kapa Biosystems library quantification kit according to manufacturer’s protocols. Uniquely indexed libraries were pooled in equimolar ratios and sequenced on Illumina NextSeq500 with single-end 75bp reads at the Dana-Farber Cancer Institute Molecular Biology Core Facilities.

### SLAM-seq

Thiol (SH)-linked alkylation for the metabolic sequencing of RNA (SLAM-seq) was performed as described (Herzog et al., 2017, Harada et al., 2022). Briefly, C2C12 cells were seeded the day before the labeling experiment that allows exponential growth (50%-80% confluency for experiment). 0.5 × 10^6^ C2C12 cells per replicate was incubated with 500 nM dTAG47/DMSO for 4h. S4U labeling was performed by adding S4U to a final concentration of 100μM for an additional hour. After labeling, cells were washed twice with 1X PBS, and lysed directly in TRIzol. Total RNA was extracted using Quick-RNA MiniPrep (Zymo Research) according to the manufacturer’s instructions except including 0.1 mM DTT to all buffers. Thiol modification was performed by 10 mM iodoacetamide treatment followed by quenching with 20 mM DTT. RNA was purified by ethanol precipitation, and RNA-seq was performed as described above.

### ATAC-seq

ATAC-seq was performed as described (Buenrostro et al., 2015). Total 50,000 cells were washed with cold PBS and lysed using cold lysis buffer (10mM Tris-HCl, pH 7.4, 10mM NaCl, 3mM MgCl2 and 0.1% IGEPAL CA-630). The pellet was then resuspended in the transposition reaction mix (25 μL 2× TD buffer, 2.5μLTn5 Transposes (Illumina) and 22.5μL nuclease-free water) and incubated at 37°C for 30min. Immediately following transposition incubation, DNA was purified using a Qiagen MinElute Kit. Transposed DNA fragments were amplified by PCR and libraries were sequenced on on Illumina NextSeq500 with pair-end reads (2X 42bp, 8bp index) at the Dana-Farber Cancer Institute Molecular Biology Core Facilities.

### ChIP-seq

ChIP-seq was performed as described with modifications (Cha et al., 2021). Briefly, ∼1 x 10^7^ cells per IP were crosslinked with 1% formaldehyde for 10 min at room temperature and followed by adding Glycine (2.5M in dH2O) to final 0.125M for 5min. Cells were washed twice with ice-cold PBS, and then scraped off the plates into 15ml Falcon tube. Nuclei were prepared using truChIP Chromatin Shearing Reagent Kit (Covaris).

Chromatin was sonicated to around 200–500 bp in shearing buffer (10mM Tris-HCl pH 7.6, 1mM EDTA, 0.1% SDS) using a Covaris E220 sonicator. The sheared chromatin was diluted and adjusted to 150mM NaCl and 1% Triton X-100, and incubated with antibody at 4 °C overnight. Protein A or G Dynabeads (Invitrogen) were added to the ChIP reactions and incubated for 3 hours at 4 °C. Subsequently, Dynabeads were washed twice with low salt wash buffer(10mM Tris-HCl PH7.4, 150mM NaCl, 1mM EDTA, 1% TritonX-100, 0.1% SDS, 0.1% sodium deoxycholate), twice with high salt wash buffer (10mM Tris-HCl PH7.4, 300mM NaCl, 1mM EDTA, 1% TritonX-100, 0.1% SDS, 0.1% sodium deoxycholate), twice with LiCl buffer (10mM Tris-HCl PH8, 1mM EDTA,0.50% sodium deoxycholate, 0.5%NP-40, 250mM LiCl) and twice with TE buffer(10mM Tris-HCl pH 8.0, 1mM EDTA, pH 8.0). The chromatin was eluted in SDS elution buffer (1% SDS, 10mM EDTA, 50mM Tris-HCl, pH 8.0) followed by reverse crosslinking at 65°C overnight. ChIP DNA were treated with RNaseA and protease K, and purified using Phenol-chloroform extraction. 2–10 ng of purified ChIP DNA was used to prepare sequencing libraries, using NEBNext Ultra DNA Library Prep Kit for Illumina (NEB) according to the manufacturer’s instructions. The finished ChIP-seq libraries were quantified by Qubit fluorometer, Agilent TapeStation 2200, and RT-qPCR using the Kapa Biosystems library quantification kit according to manufacturer’s protocols. Uniquely indexed libraries were pooled in equimolar ratios and sequenced on Illumina NextSeq500 with single-end 75bp reads at the Dana-Farber Cancer Institute Molecular Biology Core Facilities.

### CUT&RUN

CUT&RUN experiments were carried out as described (Janssens et al., 2019, Meers et al, 2019) with modifications. Briefly, ∼0.5×10^6^cells were collected and wash 3 times with Wash buffer (20 mM HEPES-KOH, pH 7.5,150mM NaCl, 0.5mM Spermidine, and Roche Complete Protease Inhibitor EDTA-free). Cells were captured with BioMagPlus Concanavalin A (Polysciences) for 10min at room temperature and incubated with primary antibody (final concentration 1:100) on a nutator at 4°C overnight. After incubation, samples were washed 3 times with Dig-wash buffer (0.025% digitonin, 20 mM HEPES-KOH, pH 7.5,150mM NaCl, 0.5mM Spermidine, and Roche Complete Protease Inhibitor EDTA-free). (Note: for primary antibodies from mouse, secondary antibody rabbit anti-mouse (final 1:100) was added at this step. Reactions were placed on the nutator at 4°C for 1h, and then washed 3 times with dig-wash buffer). Next, protein A-MNase was added at 700ng/ml in Dig-wash buffer and incubated on a nutator at 4°C for 1 hour. Samples were washed again and placed in a 0°C metal block (heating block sitting in wet ice). To activate protein A-MNase, CaCl_2_ was added to a final concentration of 2 mM. The reaction was incubated at 0°C for 30min and stopped by addition of equal volume of 2XSTOP buffer (340mM NaCl, 20 mM EDTA, 4 mM EGTA, 100ug/ml RNase A and 50ug/ml glycogen). The CUT&RUN fragments were released from the insoluble nuclear chromatin at 37°C for 30min. Digested chromatin in the supernatant were collected and then digested by proteinase K at 50°C for 1h. DNA was extracted by phenol chloroform extraction, ethanol precipitation.

### Library Preparation and Sequencing for CUT&RUN

For transcription factors (TF), libraries were constructed using the library preparation manual of “NEBNext® Ultra™ II DNA Library Prep Kit for Illumina”, NEB E7645, with modification (Liu et al., 2018). Briefly, dA-tailing temperature was decreased to 50°C to avoid DNA melting, and the reaction time was increased to 1 hour to compensate for lower enzymatic activity. After adaptor ligation, 1.87x volume of AMPure XP beads was added to the reaction to ensure high recovery efficiency of short fragments. After 12 cycles of PCR amplification (30s@98°C, 12 cycles of 10s@98°C, 10s@65°C, final extension 5min @65°C), the reaction was cleaned up with 1.2x volume of AMPure XP beads. For histone markers, libraries were prepared using NEBNext® Ultra™ II DNA Library Prep Kit for Illumina”, NEB E7645, per manufacturer’s instruction with modification. Briefly, after end repair and adapter ligation, 1.1x volume of AMPure XP beads was added to the reaction for DNA cleanup. After 12 cycles of PCR amplification (30s@98°C, 12 cycles of 10s@98°C, 10s@60°C, final extension 1min @72°C), the reaction was cleaned up with 1.1x volume of AMPure XP beads.

16-24 barcoded libraries were quantified and mixed at equal molar ratio. Library was loaded to a NextSeq 500/550 High Output Kit v2 (75 cycles), and sequenced in the NextSeq 500 platform. To enable determination of fragment length, paired-end sequencing was performed (2x42 bp, 6 bp index)

### Hi-C library preparation

Hi-C was performed as described with modification (Belaghzal et al., 2017, Golloshi et al., 2018). 5 million cells were crosslinked with 1% formaldehyde for 10 min and then quenched with glycine. Cells were lysed and then digested with DpnII overnight at 37°C. Sticky ends were filled with dNTPs containing biotin-14-dATPs at 23 °C for 4 h. Furthermore, blunt ends were ligated using T4 DNA ligase at 16 °C for 4 h. Ligation products were treated with proteinase K at 65 °C overnight to reverse cross-linking and then purified using 2 consecutive phenol-chloroform extraction. Ligation products were confirmed by agarose gel.

Biotins were removed from un-ligated ends and then fragmented to average size of 200-400 bp by sonication. Fragmented DNAs were size-selected up to 400 bp using AMPure XP beads. At this step, Hi-C libraries were prepared using adapted NEBNext protocol to work after beads pulldown (Golloshi, Rosela et al., 2018). Biotin-tagged ends were pulled down using Dynabeads MyOne Streptavidin C1 (Thermo Fisher Scientific 65001). Standard Illumina library preparation protocol including end repair, A-tailing, and adaptor ligation was performed on beads with the NEBnext Ultra II kit (New England Biolabs E7645). An optimal PCR cycle for final library amplification using NEBnext Ultra II Q5 was determined, and between six and nine PCR cycles were used in our study. Amplified PCR libraries were purified with 1.5X AMpure XP beads and were quantified by a Qubit fluorometer and Qubit dsDNA HS kit (Thermo Fisher Q32851). For quality control, a small aliquot of the final Hi-C libraries (1.5ul) was digested with ClaI enzyme at 37°C for one hour and quantified approximate% uncut DNA (correlates to the fraction of non-ligation products) from Bioanalyzer analysis. Illumina NovaSeq 50-bp paired-end sequencing (PE50) was used to obtain ∼400 million reads for each replicate.

### Data processing and analysis

#### Gene expression analysis from RNA-seq and SLAM-seq

RNA-seq experiments were processed by the HISAT pipeline, which includes alignment, filtering, and gene quantification steps. Differential expression analysis was carried out using DESeq2 using default settings. We performed SLAM-seq at 4 hrs. post dTAG47 exposure and quantified nascent RNA transcription using the SlamDunk pipeline (Neumann et al., 2019, Harada, Taku et al., 2022). Differential analysis of SLAM-seq data was performed on the transcript read counts with thymine-to-cytosine (T>C) conversions following s4U metabolic labeling (newly synthesized RNA) and also on the total transcript read counts (total RNA). DESeq2 was again used for differential nascent transcript analysis.

#### CUT&RUN data normalization and processing

We used CUT&RUNTools (Zhu et al., 2019) to process MyoD, YY1, CTCF, and Rad21 CUT&RUN experiments from raw fastq reads. This process included reads trimming, alignment to the reference genome (mm10), BAM file duplicate marking, and fragment filtering based on fragment size. Next, we performed background-based data normalization on the group of experiments to be normalized together (i.e. Matr3 dTAG47 & DMSO, or Matr3 WT & KO of a given antibody). For experiments per group, we pooled all experiments’ BAM files and performed MACS2 (Zhang et al., 2008) peak calling to get N peaks. Peak-flanking regions were obtained, and stored for the purpose of normalization. Bamliquidator was called to compute the number of reads in each peak-flanking region in each experiment. DESeq2’s estimateSizeFactor() (Love et al., 2014) was called on the matrix of peak-flanking region reads over all experiments to calculate a scale factor per experiment. With the scale factor computed, the original BAM file was subsampled at a rate equal to the scale factor to get a normalized BAM file. Finally, MACS2 was called to generate peaks from the normalized BAM file. A normalized bigwig file was used for visualization.

#### ATAC-seq data normalization and processing

ATAC-seq data were processed using a procedure similar to CUT&RUN described above, which included reads trimming, alignment to the reference genome, BAM duplicate marking, and fragment size filtering. A special note is that we kept paired-end fragments that are <150bp instead of <120bp setting for TF CUT&RUN. For ATAC-seq data normalization, we adopted the same background-based data normalization as used in CUT&RUN (see CUT&RUN section).

#### Differential binding analysis of CUT&RUN and ATAC-seq data

To identify differential binding between dTAG47 & DMSO or WT & KO, experiments to be compared were first normalized using background-based data normalization (see above). Next, we pooled the BAM files and called peaks using MACS2 to generate a set of common regions on which to test differential binding. Bamliquidator was used to compute the number of reads in each common region in each experiment. Finally, DESeq2 was called to identify regions that exhibit differential binding.

#### ChIP-seq analysis

For analysis of CTCF and Rad21 ChIP-seq, we used a ChIP-seq data processing pipeline to perform reads trimming, alignment to the reference genome (mm10), and BAM file duplicate marking. Next, we performed background-based data normalization on the group of experiments to be normalized together (i.e. Matr3 dTAG47 & DMSO, or Matr3 WT & KO of a given antibody), similar to the normalization procedure used in CUT&RUN data. Once samples were normalized, we proceeded to call peaks using MACS2, and then differential binding analysis using DESeq2, same as described in the CUT&RUN paragraph.

#### Gene set enrichment analysis

We mapped differential ATAC-seq peaks to the nearest genes that are 25kb from transcription start site (TSS). Each differential peak contributed a score to the gene that equals to the 1/n * (−10logP) where n is the number of genes within 25kb vicinity of the peak, and P is the P-value significance of the differential peak. We next input the top scoring genes to DAVID Functional Annotation Tool (https://david.ncifcrf.gov/tools.jsp) for gene enrichment analysis using default settings. To ensure the robustness of the gene enrichment results, we also performed enrichment using SEEK (https://seek.princeton.edu). Here the top scoring genes were input as a query and co-expressed genes to the query were identified and were used to perform enrichment analysis. We used the default settings (top 500 co-expressed genes and GO: biological process).

#### Hi-C processing and analysis

We used HiC-Pro (Servant et al., 2015) to align reads to mm10 reference genome assembly. By providing the DpnII restriction enzyme digestion sites and upon further filtering, HiC-Pro returns valid interaction pairs for each of four conditions: dTAG47, DMSO, WT, and KO. Next, we used the HIFI tool (Cameron et al., 2020) to impute interactions for our Hi-C data based on the observed over expected interaction frequency matrix. The interaction frequency matrix was at restriction fragment resolution, roughly equivalent to ∼5kb resolution between contact sites. HIFI employs a Markov random field-based imputation to fill in missing values, and smooth neighboring values in the contact matrix to enhance loop detection. Initial evaluation found that this additional imputation step improved the detection of loops from Hi-C data without loss of resolution and increased the agreement between different loop calling methods (Supplemental Figure 8). We next performed loop calling on the imputed interaction frequency matrix using the Hiccup similar procedure that is implemented within HIFI. For differential loop analysis, we had three Hi-C replicates per DMSO and dTAG47 groups. After calling loops in individual replicate samples, we pooled loops to get a union of interaction regions on which to do differential analysis. For each (i,j) interaction pair, we went back to each sample to compute the Hiccup (Rao et al., 2014) score in each sample which was defined by: score(i,j) =P(i,j)/[max(D(i,j), H(i,j), V(i,j), BL(i,j)], where P(i,j) is the average interaction frequency (IF) in the peak. D(i,j), H(i,j), V(i,j), and BL(i,j) are several types of flanking regions to which P(i,j) is compared against.

Hiccup scores were z-score normalized within each sample across all pairs. Differential loop scores were obtained by: zDmso1(i,j) - zDtag1(i,j), zDmso2(i,j) - zDtag2(i,j), zDmso3(i,j) - zDtag3(i,j), …, for all combinations of DMSO and dTAG replicates, resulting in a set of difference scores DS. A final meta-differential loop score Z(i,j) was returned by Stouffer z-score meta-analysis method: Z(i,j) = sum_k(zDmsox(i,j) - zDtagx(i,j)) / sqrt(k) where k=|DS|. Z(i,j) that correspond to P-value <0.05 are deemed as significant. For compartment analysis, A/B compartments were identified using the “runHiCpca.pl” script in HOMER (cite PMID:30146161) at the resolution of 50kb. The “+/-” sign of the compartment was determined by TSS enrichments, where compartments showing enrichment of TSS were assigned the ‘+’ sign and those showing depletion of TSS were assigned the ‘-’ sign. To visualize the changes in compartmentalization strength, we generated and compared the saddle plots. First, for each chromosome, we removed the 1% genomic bins with the lowest sequencing coverage in order to remove the bias caused by insufficient coverage. Then we ranked the remaining genomic bins by the PC1 scores from high to low. We reordered the rows and columns of the contact matrix according to the same ordering. Then the contact map was coarse-grained into a 100*100 matrix, where the element (m, n) represents the mean interaction frequency between bins of the m-th percentile and the n-th percentile. The average of the coarse-grained contact matrices from all chromosomes were then plotted as the saddle plot. The obs/exp contact matrices at 50kb were used for this analysis. To quantify the intra- and inter-compartment interaction strength, the average interaction strength for genomic bins with PC1 values in top 25% and bottom 25% are used to measure the AA (top left), BB (bottom right) and AB (top right) interaction strengths. To visualize the differences in saddle plots of two samples, we plotted the fold change of the two contact matrices in an element-wise way after log2 transformation

#### Visualization of CUT&RUN heatmaps and Hi-C interactions

To generate comparative heatmaps between CUT&RUN groups (DMSO vs dTAG47), we used Deeptools’s computeMatrix and plotHeatmap functions on the list of differential binding sites. For visualizing Hi-C interaction matrix, we used the HIFI visualization capability to generate a fragment-resolution interaction heatmap. To interactively navigate the Hi-C data, we used the Washington University Epigenome Browser (http://epigenomegateway.wustl.edu/). We prepared the interaction data in the tabix and longrange format as required by the browser, and loaded the tracks. We chose the “arc” representation to visualize locus-specific interactions. The “heatmap” mode was chosen to visualize all interactions in a given region, omitting interactions that are outside, and visualizing gained and lost interactions in shades of greens and reds respectively.

