## Supplementary figures and images for "Matrin3 mediates differentiation through stabilizing chromatin accessibility and chromatin loop-domain interactions, and YY1 mediated enhancer-promoter interactions"

### Supplemental Figure 1

Supplemental Figure 1

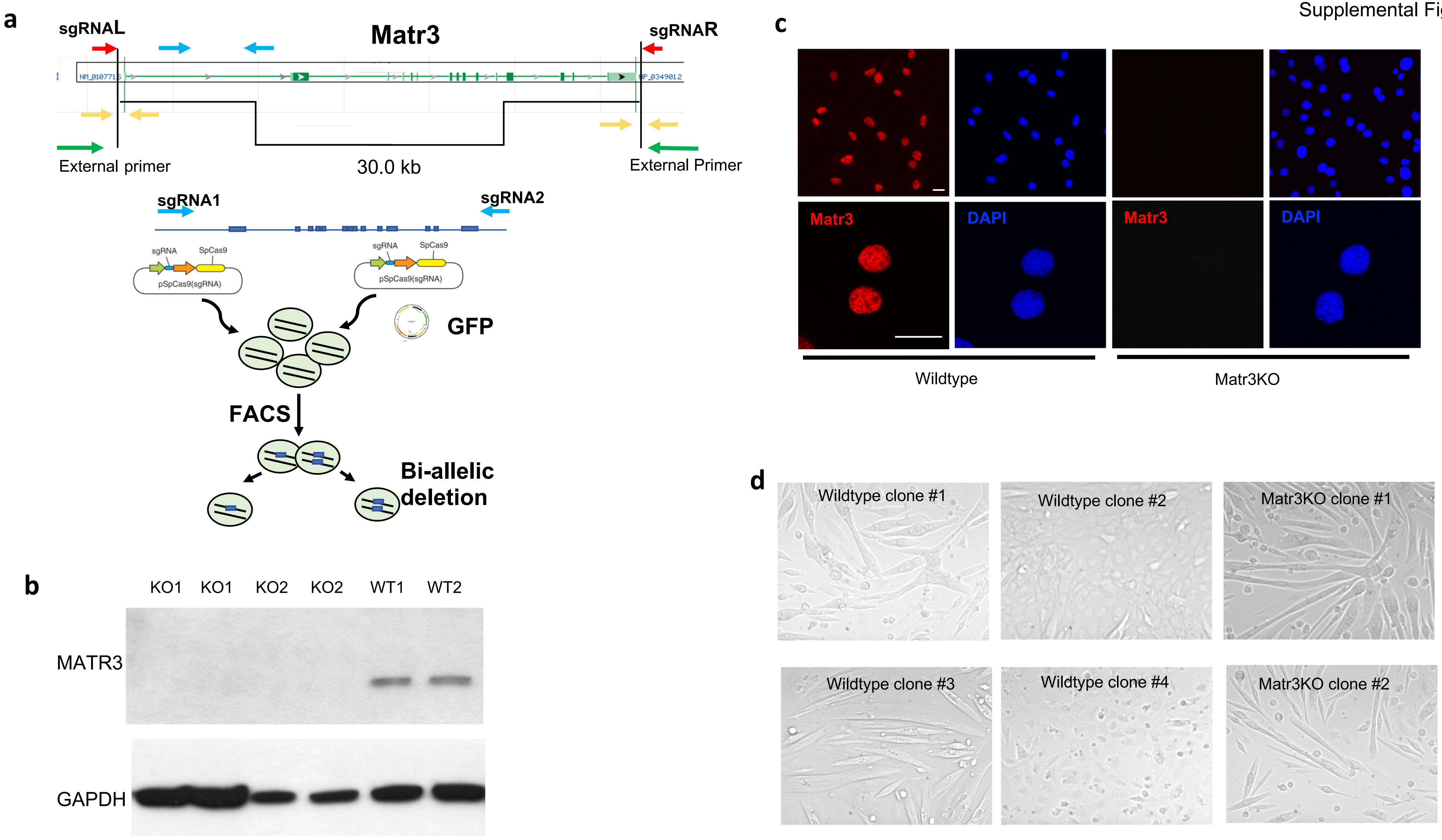

### Supplemental Figure 2

Supplemental Figure 2

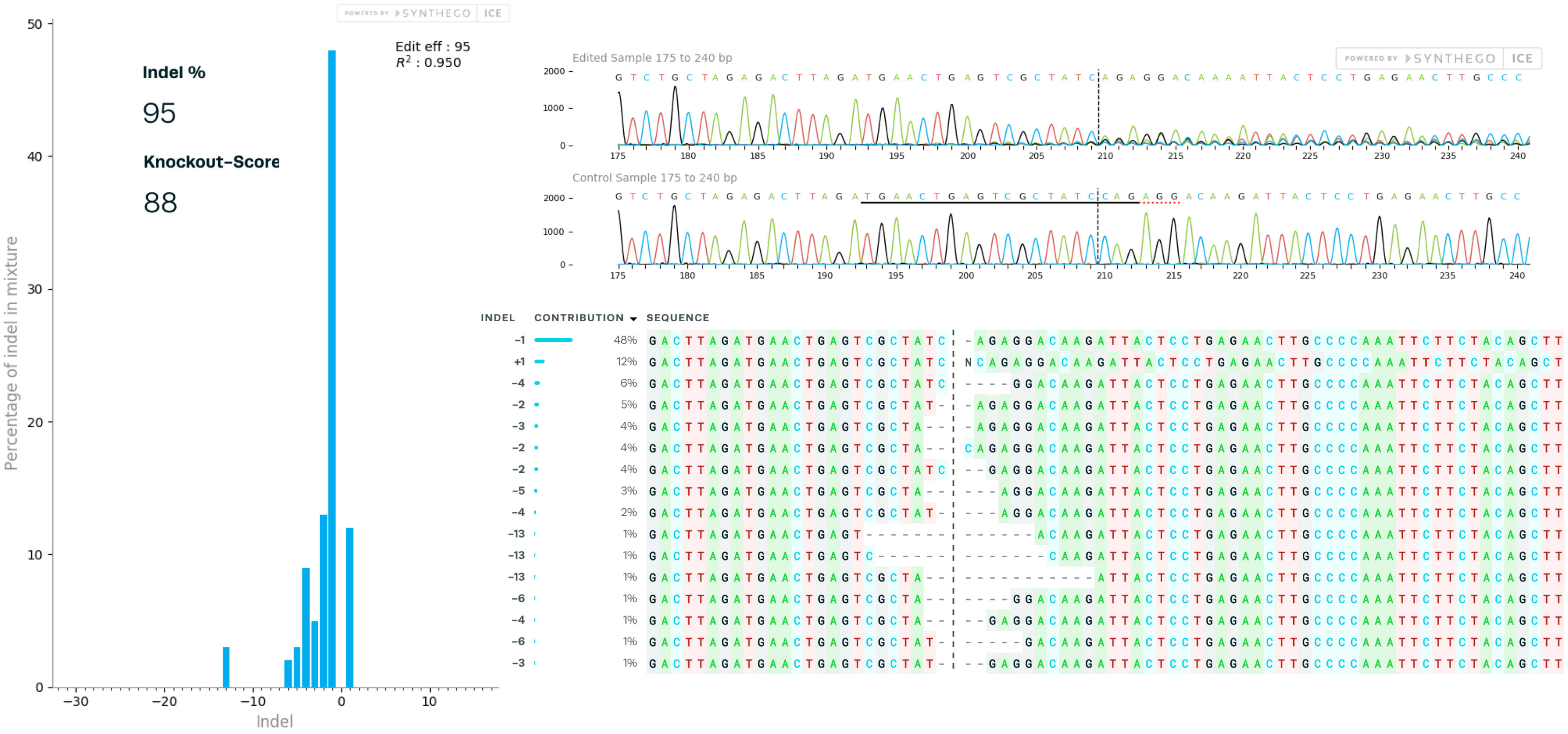

### Supplemental Figure 3

Supplemental Figure 3

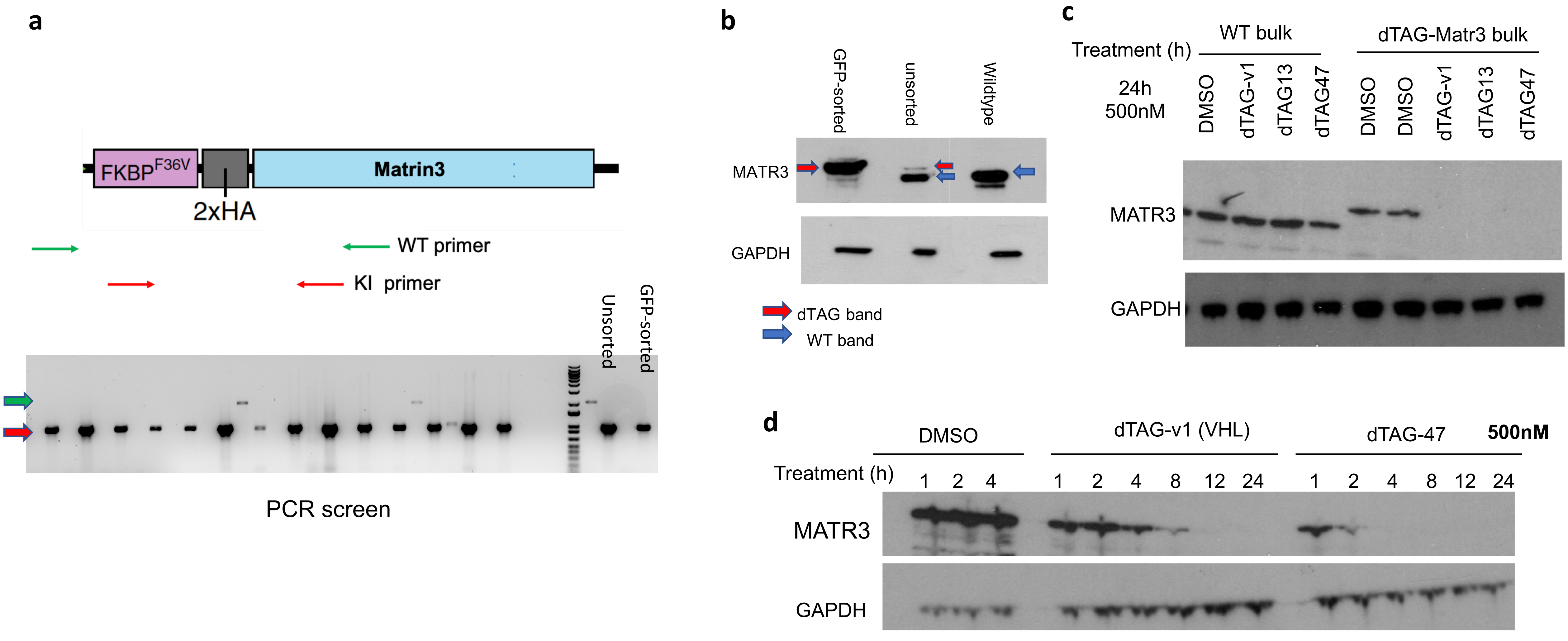

### Supplemental Figure 4

#### Supplemental Figure 4

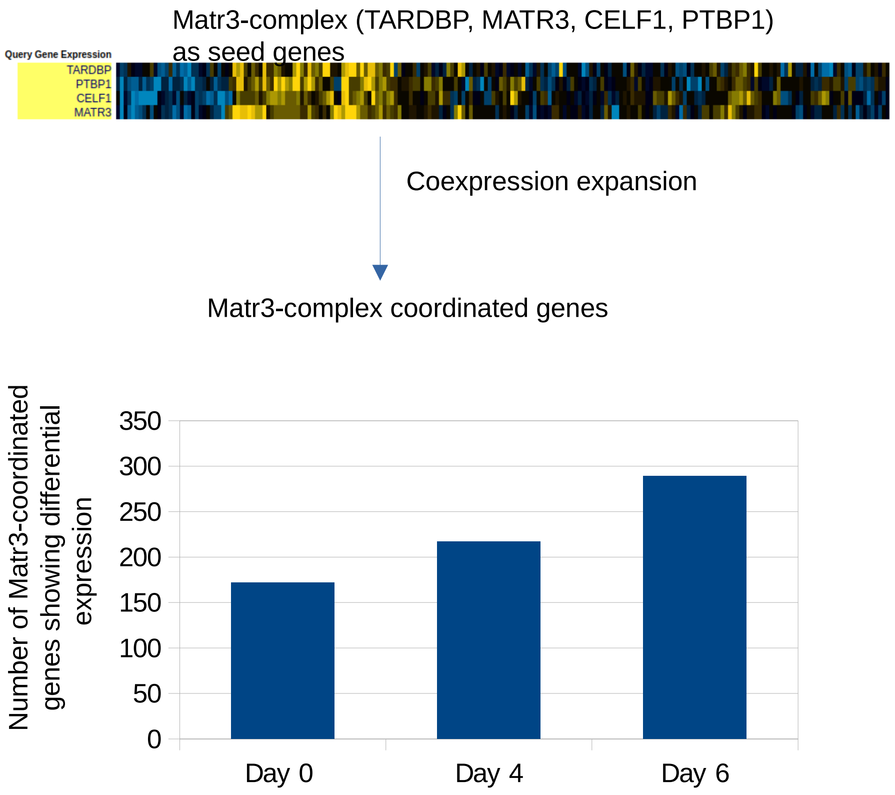

### Supplemental Figure 5

Supplemental Figure 5

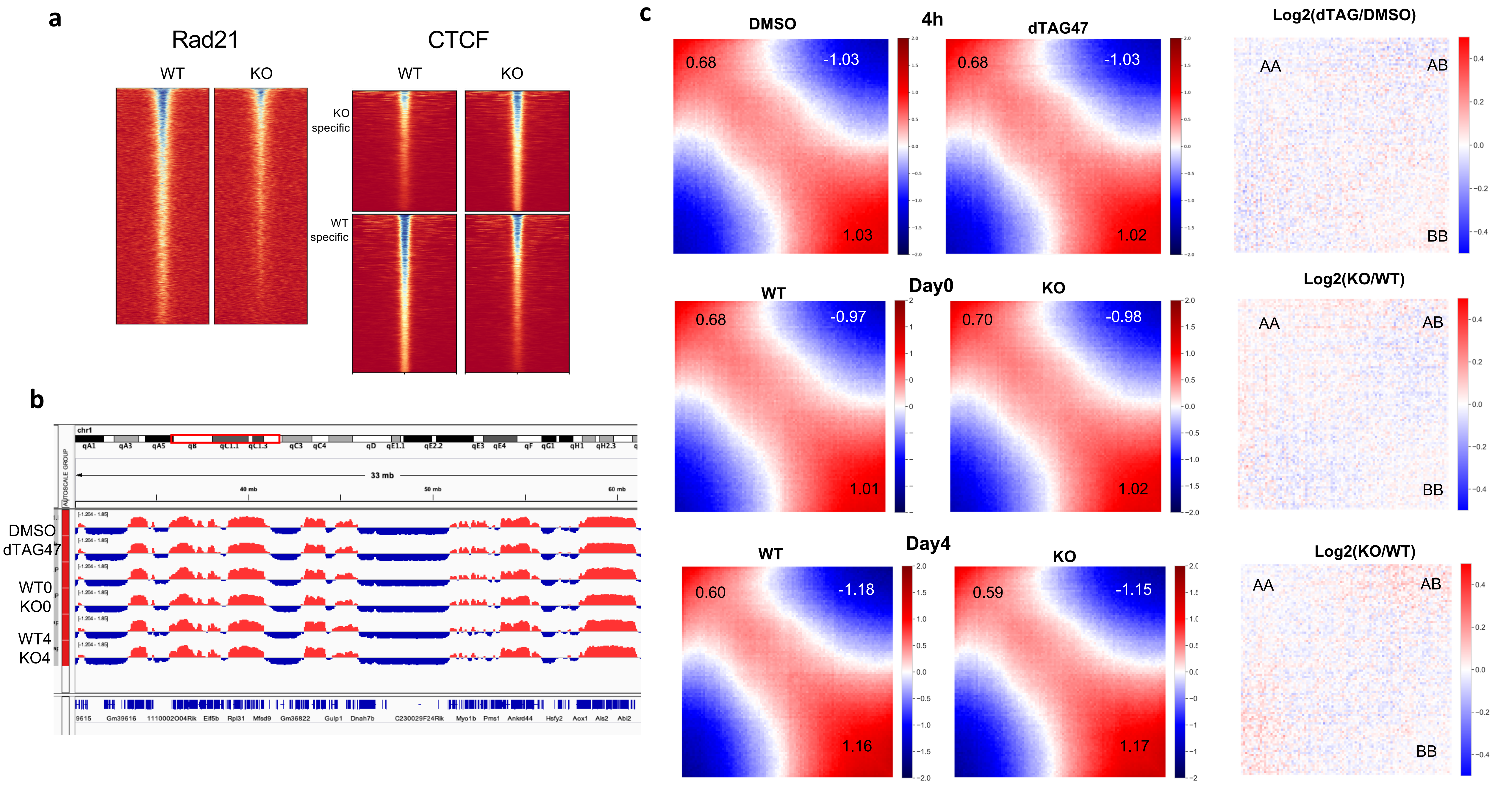

### Supplemental Figure 6

Supplemental Figure 6

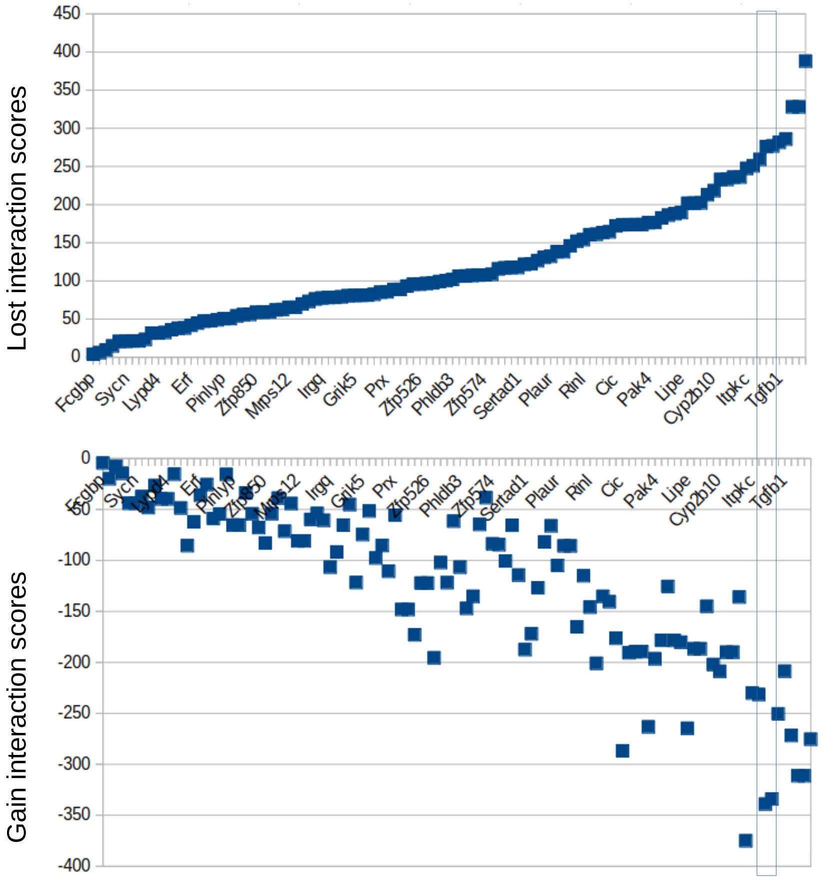

### Supplemental Figure 7

Supplemental Figure 7

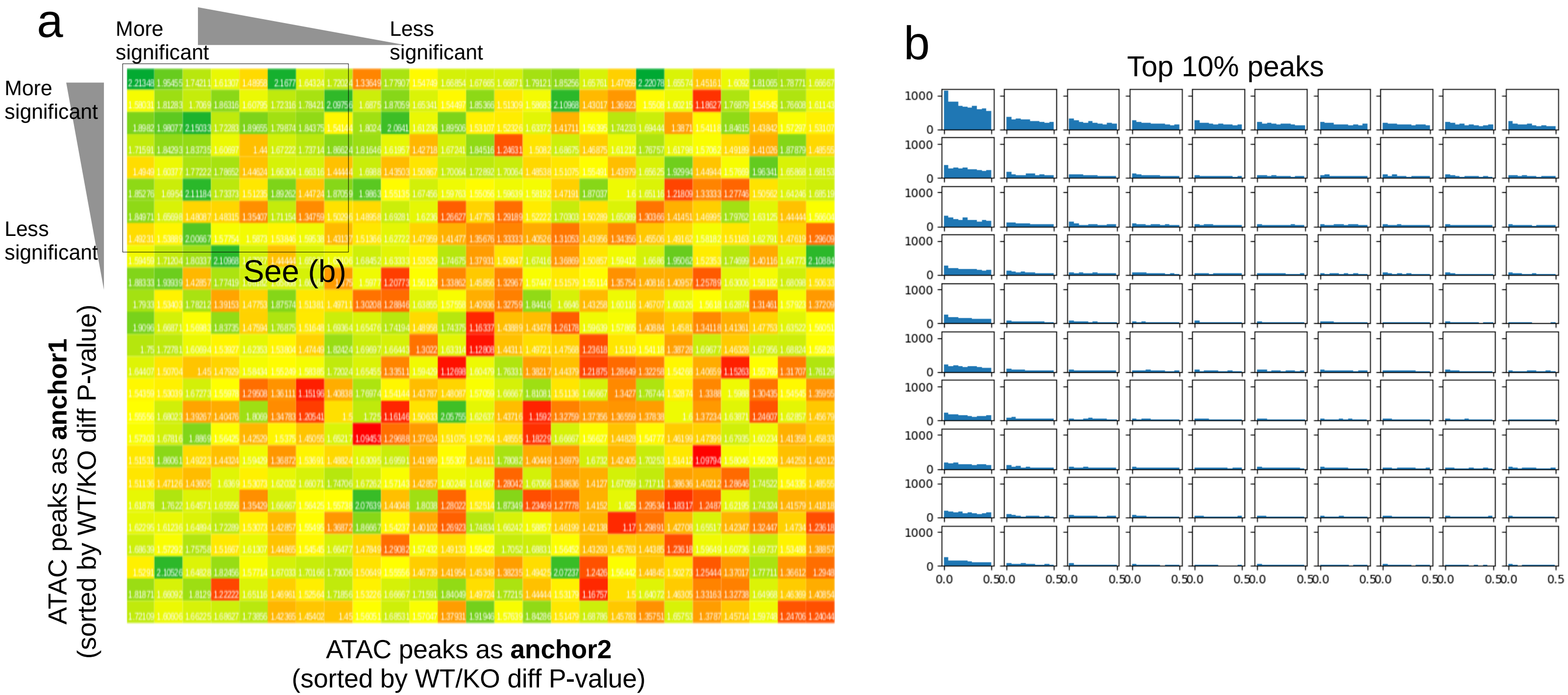
