## Supplemental Figure 8 for "Matrin3 mediates differentiation through stabilizing chromatin accessibility and chromatin loop-domain interactions, and YY1 mediated enhancer-promoter interactions"

**a** Loop calling at various resolution with Hiccup algorithm

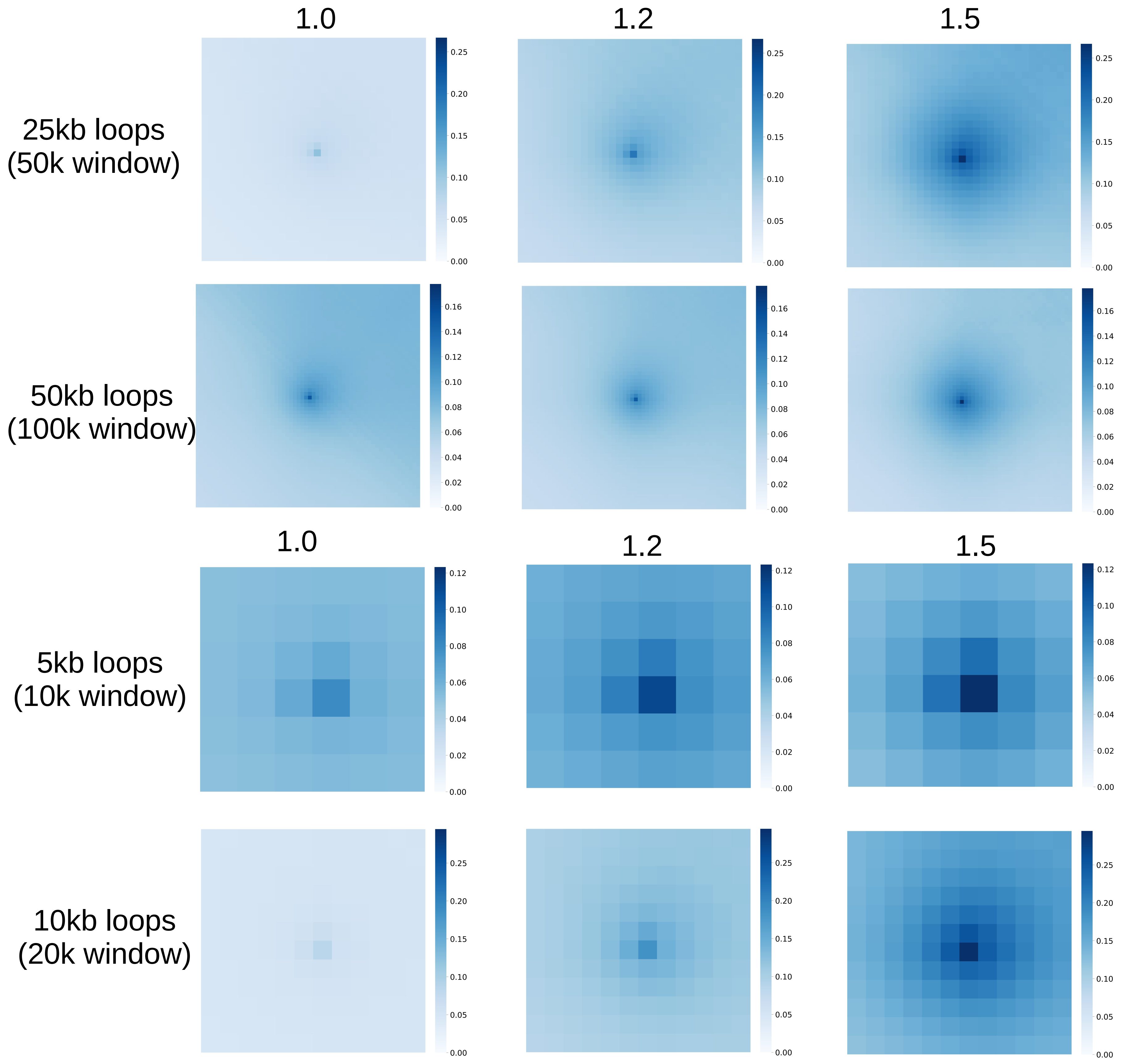


**b** Loop calling comparison, agreement between Hiccup and FitHiC with or without HIF imputation

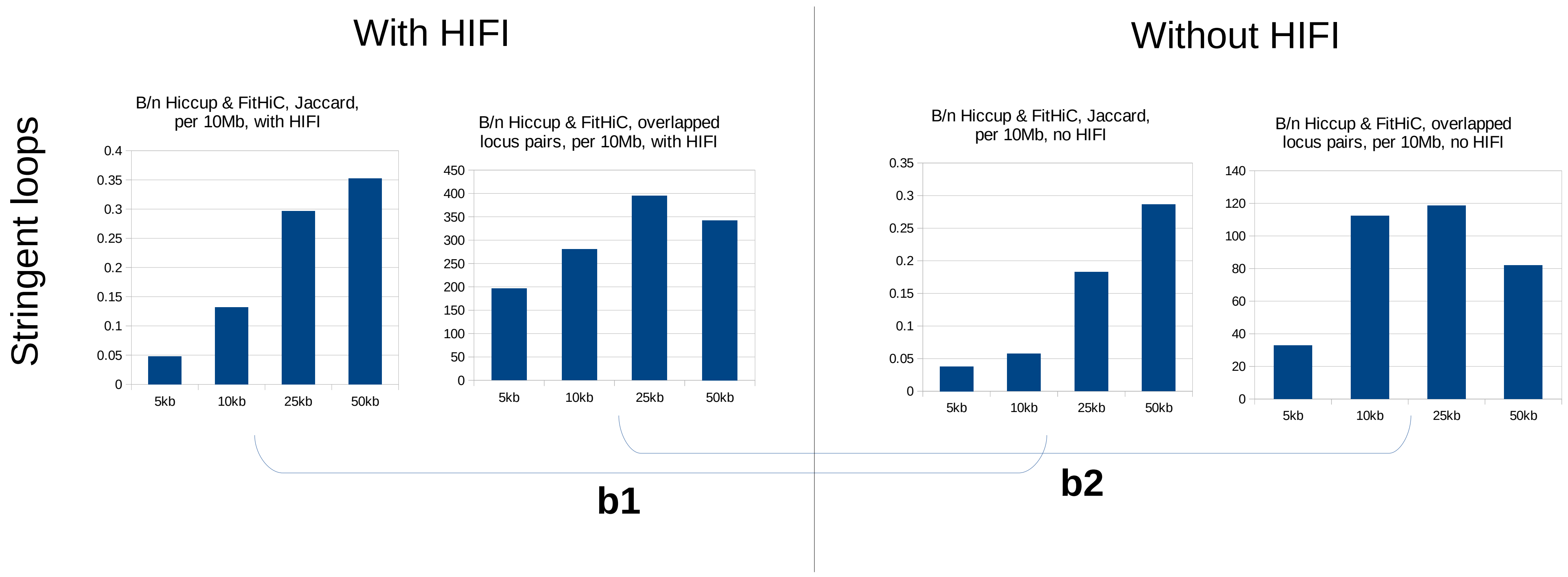
